# Inhibition underlies fast undulatory locomotion in *C. elegans*

**DOI:** 10.1101/2020.06.10.138578

**Authors:** Lan Deng, Jack Denham, Charu Arya, Omer Yuval, Netta Cohen, Gal Haspel

**Affiliations:** Jersey Institute of Technology; University of Leeds

## Abstract

Inhibition plays important roles in modulating the neural activities of sensory and motor systems at different levels from synapses to brain regions. To achieve coordinated movement, motor systems produce alternating contraction of antagonist muscles, whether along the body axis or within and among limbs. In the nematode *C. elegans*, a small network involving excitatory cholinergic and inhibitory GABAergic motoneurons generates the dorsoventral alternation of body-wall muscles that supports undulatory locomotion. Inhibition has been suggested to be necessary for backward undulation because mutants that are defective in GABA transmission exhibit a shrinking phenotype in response to a harsh touch to the head, whereas wild-type animals produce a backward escape response. Here, we demonstrate that the shrinking phenotype is exhibited by wild-type as well as mutant animals in response to harsh touch to the head or tail, but only GABA transmission mutants show slow locomotion after stimulation. Impairment of GABA transmission, either genetically or optogenetically, induces lower undulation frequency and lower translocation speed during crawling and swimming in both directions. The activity patterns of GABAergic motoneurons are different during low and high undulation frequencies. During low undulation frequency, GABAergic VD and DD motoneurons show similar activity patterns, while during high undulation frequency, their activity alternates. The experimental results suggest at least three non-mutually exclusive roles for inhibition that could underlie fast undulatory locomotion in *C. elegans*, which we tested with computational models: cross-inhibition or disinhibition of body-wall muscles, or inhibitory reset.

**Significance Statement:** Inhibition serves multiple roles in the generation, maintenance, and modulation of the locomotive program and supports the alternating activation of antagonistic muscles. When the locomotor frequency increases, more inhibition is required. To better understand the role of inhibition in locomotion, we used *C. elegans* as an animal model, and challenged a prevalent hypothesis that cross-inhibition supports the dorsoventral alternation. We find that inhibition is related to the speed rather than the direction of locomotion and demonstrate that inhibition is unnecessary for muscle alternation during slow undulation in either direction but crucial to sustain rapid dorsoventral alternation. We combined calcium imaging of motoneurons and muscle with computational models to test hypotheses for the role of inhibition in locomotion.

## Introduction

Alternating activation of antagonistic muscles is ubiquitous in motor programs. During locomotion, limbed animals alternate flexor–extensor muscles as well as their left and right counterparts, while limbless locomotion involves antagonistic axial muscles that generate body bending. The generation of an alternating pattern often involves direct or indirect inhibitory pathways to the antagonistic motoneuron pool (Kiehn, 2011, 2016). Inhibition serves multiple roles in the generation, maintenance, and modulation of the locomotive program. Hypotheses range from cross-inhibitory induced alternation of antagonists to postinhibitory rebound contributing to rhythm generation. Ascending interneurons produce recurrent inhibition of sensory pathways that gate reflex responses and limit firing of central pattern generator (CPG) neurons; while descending interneurons can produce tonic or phasic inhibition that reduces responsiveness and spontaneous locomotion, or stops it all together (Roberts *et al*., 2008). In particular, premotor inhibition is necessary for rapid locomotion in insects (Gowda *et al*., 2018 Kohsaka *et al*., 2014), amphibians (Li and Moult, 2012), fish (Liao and Fetcho, 2008; Satou *et al*., 2009) and mammals (Gosgnach *et al*., 2006; Kiehn, 2016; Zhang *et al*., 2014). Across fields very disparate from neuroscience, through ecological and genetic networks, to pedestrian and vehicle traffic, negative regulation plays a role in modulating and coordinating the performance of system-parts to overall speed up processes of complex systems (Gershenson and Helbing, 2015). For example, negative autoregulation of gene networks, in which a gene product plays as a repressor of their transcription, speeds up the response time of gene circuits, while also promoting robustness (Alon, 2007).

In *C. elegans*, GABAergic motoneurons in the ventral nerve cord have been suggested, based on their morphology, connectivity, and mutant phenotype, to provide cross-inhibition necessary for backward crawling (McIntire *et al*., 1993a; McIntire *et al*., 1993b; Schuske *et al*., 2004). The locomotion circuit is composed of 75 motoneurons, divided by their morphology into 6 excitatory cholinergic and 2 inhibitory GABAergic classes that innervate 95 body-wall muscles. The motoneurons synapse onto thin processes called muscle arms that extend from muscle cells onto the ventral cord (White *et al*., 1976, 1986; Chen *et al*., 2006; Emmons, 2015; Altun and Hall, 2009). Muscle arms are functionally analogous to vertebrate motoneurons, while *C. elegans* motoneurons are functionally analogous to vertebrate spinal interneurons. GABAergic inhibitory motoneurons (namely 6 DD and 13 VD motoneurons that innervate dorsal and ventral muscles, respectively) receive inputs only from other motoneurons (White *et al*., 1976, 1986; Haspel and O’Donovan, 2011; Petrash *et al*., 2013). Each VD receives input from the dorsal cholinergic motoneurons (DA, DB, and AS) that innervate the opposing muscle cells, while DD motoneurons receive input from the ventral cholinergic motoneurons (VA and VB) (White *et al*., 1976, 1986; Haspel and O’Donovan, 2011). In addition to their synaptic output at neuromuscular junctions, opposing DD and VD motoneurons also innervate each other, whereas VD motoneurons also innervate the local VA and VB (White *et al*., 1986; Haspel and O’Donovan, 2011).

Mutant strains of *C. elegans* that are defective in any part of their GABAergic transmission (the so-called *shrinker* mutants) respond to anterior noxious stimuli, such as a harsh touch to the head, with coactivation instead of alternation of antagonistic dorsoventral muscle, producing a shrinking response instead of the rapid reverse undulation seen in the wild type. The mutant animals as well as animals with laser-ablated GABAergic motoneurons, shorten the anterior portion of their body and have been reported not to move backward while performing normal forward crawling, albeit with decreased undulatory amplitude (McIntire *et al*., 1993a; McIntire *et al*., 1993b). Hence, it was suggested that GABAergic motoneurons mediate dorsoventral cross-inhibition during backward locomotion (McIntire *et al*., 1993b), or direction change (Schuske *et al*., 2004). According to this hypothesis, when an excitatory cholinergic A motoneuron (e.g., VA) activates postsynaptic muscle arms, it also activates an inhibitory GABAergic motoneuron (e.g., DD) that prevents excitation of the opposing muscle; shrinking may be caused by the coactivation of dorsal and ventral A-type (DA and VA) motoneurons, or by coactivation by A- and B-type (DB and VB) motoneurons. An alternative hypothesis suggests that neural, rather than muscular inhibition, plays an important role (Boyle *et al*., 2012). In a computational model testing this hypothesis, abolishing neural inhibition from VD to VB is detrimental to rapid but not to slower undulation.

Here, to assess the relative contributions of these hypotheses, we quantified the response to harsh touch and spontaneous crawling and swimming behaviors of different GABA transmission mutants as well as an optogenetic strain in which GABAergic motoneurons can be inactivated acutely. We recorded the activity of GABAergic motoneurons with calcium imaging during low and high undulation frequencies. Finally, we suggested competing hypotheses for the roles of inhibition in *C. elegans* locomotion and tested them with computational models.

## Methods

### C. elegans strains

We maintained *C. elegans* strains under standard laboratory conditions on Nematode Growth Medium (NGM) agar plates with OP-50-1 *Escherichia col*i (*E. coli*) bacterial lawn at 20°C (Brenner, 1974; Corsi *et al*., 2015). All animals used in the experiments were hermaphrodites.

The *C. elegans* strains used in this study were N2 (wild type), CB120, CB1562, CZ1632, TM5487, TM5916, TOL11, TOL12, TOL15, and VC1433. We obtained N2 (a reference laboratory strain considered as wild type), VC1433 *[unc-25 (ok1901) III]*, CB1562 *[vab-7(e1562) III]*, CB120 *[unc-4(e120) II]*, and CZ1632 (*juIs76 [unc-25p::GFP + lin-15(+)]II*) strains from the Caenorhabditis Genetic Center of University of Minnesota (https://cgc.umn.edu/), and TM5916 *[unc-46 (tm5916)]V* and TM5487 *[unc-49 (tm5487)]III* from the National Bioresource Project of Japan (https://shigen.nig.ac.jp/c.elegans/top.xhtml). We generated TOL11 (*aatIs11[myo-3p::R-CaMP2; unc-25p::G-CaMP6; unc-47p::ChR2::P2A::DsRed2; odr-1p::DsRed]III* and TOL12 (*aatIs12[unc-47p::Arch::GFP, myo-2p::mCherry*]) by integrating the extrachromosomal arrays in *jqEx456 [myo-3p::R-CaMP2; unc-25p::G-CaMP6; unc-47p::ChR2::P2A::DsRed2; odr-1p::DsRed](Inoue et al*., *2015)*, and OKA074 (*ncEx2351 [unc-47p::Arch::gfp, myo-2p::mCherry]*) (Okazaki *et al*., 2012) into the animals’ genome via UV irradiation methods, respectively; then we back-crossed the integrated strains with wild type N2 for at least 5 generations and screened for the homozygous (Ahringer, 2006). We generated TOL15 by crossing TOL14 (*aatIs4 [myo3p::nls::RCaMP2]V*) and ZW495 (*zwIs132 [myo-3p::GCamp2 + lin-15(+)]*) and screening for homozygotes for *aatIs4* and *zwIs132*.

### Free locomotion behavioral tracking

We recorded the free crawling and swimming of wild type, GABA transmission knockouts, and an optogenetic strain on NGM agar plates or in NGM buffer solution using a static multi-worm tracker. The tracker was composed of three major parts. From top to bottom, a camera (Basler ace acA4024-29um) connected to a fixed focal length lens (C Series 5MP 35mm 2/3” fixed focal length lens, Edmund Optics) with an infrared cut-off filter (M25.5×0.5 Mounted IR Cut-Off Filter, Edmund Optics), a specimen stage for plates or slides, and an infrared LED light (M850L3, Thorlabs) mounted with a collimation adaptor (COP1-B-Collimation Adaptor, Thorlabs).

One day before recording, we transferred fourth larval stage (L4) animals onto new NGM plates with OP-50-1 *E. coli* bacterial lawn so that experimental animals would be first-day young adults. During tracking, at least 30 animals were freely moving on 35 mm NGM plates without bacterial lawn or in NGM solution between a microscope slide and a coverslip, 1 mm apart. We recorded at least 9 videos with Pylon Viewer (Basler pylon Camera Software Suite) at 25 frames per second (fps) for 15 to 20 s. During video acquisition, we cropped the video frame dimensions to approximately 2,000×2,000 pixels.

One day before behavioral tracking of the optogenetic strain TOL12 (expressing Archaerhodopsin-3 (Arch, Chow *et al*., 2010) in GABAergic neurons), we transferred L4 animals onto NGM plates that had been freshly seeded with 50 mM all-trans-retinal (ATR) in OP-50-1 *E. coli*. To allow the animals to incorporate ATR, we kept the plates in the dark at 20°C for 24 h. Animals fed with ATR but recorded in the dark (under infrared LED) were tracked as a negative control group, and the same animals under lime-color LED light that activated Archaerhodopsin-3 were the experimental group. As another negative control group, TOL12 animals that were not fed with ATR were not affected by the lime-color activation light of the optogenetic protein during crawling or swimming (n=30 animals of 3 plates, data not shown). We used lime-color light (M565L3, 565 nm LED with a Collimator), which is the optimal activation light of Archearhodopsin-3 (Mattis *et al*., 2011; Okazaki *et al*., 2012), with a working illumination power of 10 mM/cm^2^. To prevent desensitization of TOL12 to the activation light, the maximum time of light exposure was 1 m (Okazaki *et al*., 2012). The rest of the operations and software settings were the same as the behavior tracking aforementioned.

### Head and tail harsh touch experiments

We transferred L4 animals to new OP-50-1 *E. coli*-seeded NGM plates one day before the experiment so that all assays were done on young adult hermaphrodite animals. During the experiment, we transferred animals onto 60 mm NGM plates without bacterial lawns. We touched the head or tail of the animals using a blunt glass probe while they were crawling forward. We acquired the videos continuously with a CMOS camera (U3CMOS14000KPA, ToupTek) and acquisition software (ToupView, https://www.touptek.com/) at 30 fps and 1,048×822 pixels greyscale during several stimulations of different animals. We further trimmed these videos into clips with one stimulation each by video editing softwares Premiere Pro (Adobe) or Shotcut (https://shotcut.org/). We also tested the response of wild-type young adult hermaphrodites to a weak chemical repellant: 0.1% sodium dodecyl phosphate (SDS) off-food, and a strong chemical repellant: 50 mM copper sulfate (CuSO_4_) on a thin layer of food (Hilliard *et al*., 2004; Ezcurra *et al*., 2011; Hart, 2006). To make sure that harsh touch does not mechanically cause changes in body length we paralyzed young adult animals with 1 mM Tetramisole hydrochloride (Sigma-Aldrich, 10 m immersion). We touched the head or tail of paralyzed animals using a blunt glass probe, acquired videos, and analyzed body length as described above. There was no significant change in body length of paralyzed animals (99 ± 1%; p=0.21 paired T-test, Table 2-1).

### Behavioral tracking analysis

We processed crawling and swimming tracking using Tierpsy Tracker v1.4.0, a multi-worm behavior tracker software (https://github.com/ver228/tierpsy-tracker/releases/) (Javer *et al*., 2018). Then, we analyzed the processed results in hdf5 files using custom programs (Code 1-1) written in Matlab (Mathworks).

Each video had between 10 and 30 animals each assigned with an index by Tierpsy. We analyzed the translocation speed and undulation frequency during forward and backward locomotion from the midbody of the animals as well as primary wavelength and maximal amplitude. Tierpsy computes the undulation frequency using the waveform frequency from the largest peak via the Fourier transform over a time window.

We used Tierpsy Tracker v1.4.0 to extract the aforementioned kinetic parameters from trimmed video clips before and after head or tail stimulation. We calculated the means of these 4 parameters within 5 s before and 5 s after the stimulation from trimmed videos of head and tail stimulation experiments.

### Changes in body length

We analyzed the body length at different time in the trimmed clips in each stimulation direction of N2 strain and each of the GABA transmission knockout strains, as well as head stimulation of *vab-7* mutant animals (CB1562) and tail stimulation of *unc-4* mutant animals (CB120): before the harsh stimulation, immediately after the stimulation, as well as 0.5 s, 1 s, and 2 s after the stimulation. We used ‘Freehand Line’ tool in ImageJ (https://imagej.nih.gov/ij/) to measure the body length of animals. We expressed the body length as the percentage of pre-stimulus length and plotted the means of normalized body lengths for each strain at each time point.

### Calcium imaging in a microfluidic device

To repeatedly measure activity at particular phases of the locomotor cycle, we placed young adult animals in a waveform silicon microfluidic device with channels designed to restrict their motion to a pattern that mimics undulatory locomotion. The cycle was fixed in space and predictable at any location along the path. The wavelength, amplitude, and width of the channel were 686.5, 322.5, and 133.5 µm, respectively. To unify the frame of reference, we arbitrarily designated the phase of a maximal dorsal bend as 90° and a maximal ventral bend as 270° and defined the rest of the phase positions accordingly. We presented collated recordings during forward locomotion from motoneurons and body-wall muscle with phase (rather than time) being the common frame of reference.

We transferred healthy fourth larval stage (L4) animals to new plates seeded with OP-50-1 *E. coli* one day before imaging. We added drops of methyl cellulose solution onto the face of a microfluidic device, and then transferred the animals into these drops. To seal the microfluidic device, we flipped covered it with an 18 mm diameter round coverslip, flipped the coverslip so the device is on top for the inverted microscope and sealed by pressing on the device. The transgenic animals could crawl freely inside the channels, and we manipulated their undulation frequency by changing the ambient viscosity. We used 3% and 0.5% (w/w) methyl cellulose in NGM buffer while recording calcium level changes in GABAergic motoneurons in TOL11, 1.5% (w/w) methyl cellulose to record calcium level changes in body-wall muscles in TOL15, and 3% and 0.5% (w/w) methyl cellulose to image GFP (as a negative control calcium-insensitive fluorescent protein to determine motion artifact) in GABAergic motoneurons in CZ1632. We only recorded calcium level changes in animals that moved smoothly in the microfluidic channel.

To image and record the fluorescence change of GCaMP or GFP, we used an inverted microscope (Olympus IX73) with a 40x/NA0.95 UPlanSApo objective and an sCMOS camera (Hamamatsu Orca Flash 4.0) and a solid-state excitation light source (X-Cite 120LED) with a dual-band filter set (Chroma 59022 ET-EGFP/mCherry), and a dual camera image splitter (TwinCam, Cairn Research with a T565lpxr dichroic beam splitter). We used Micro-Manager (v1.4; https://micro-manager.org/) to acquire multiple videos of animals moving through different parts of the channel (i.e., different locomotor phases). We saved videos as separate tiff files (2,048×2,048 pixel resolution) at a frame rate of 100 fps with 10 ms exposure for animals crawling in 0.5% (w/w) methyl cellulose, and 50 fps with 20 ms exposure for animals crawling in 1.5% (w/w) and 3% (w/w) methyl cellulose. We did not use NGM with less than 0.5% (w/w) methyl cellulose because it resulted in discontinuous and less smooth locomotion. Between acquisitions, we moved the microscope stage manually to follow animals when they moved out of view. We recorded activity in muscle cells and motoneurons in the mid-body: dorsal muscle cells 9-21 (n=24), ventral muscle cells 9-21 (n=27), DD motoneurons 2-5 (n=26), VD motoneurons 3-10 (n=59).

We used a customized Matlab program (Code 7-1) to analyze the change in fluorescence intensity of identifiable body-wall muscle cells or somata of GABAergic motoneurons. For each frame, we measured the mean fluorescence intensity of the top 50% pixels within the manually traced region of interest (ROI) around the cell of interest (F_top50%_). We then calculated and subtracted the median intensity of the frame as the background (F_BG_). Finally, we demeaned the value by dividing it by the mean fluorescence signal of this cell in the video of n frames: ΔF/F = (F_top50%_-F_BG_) / (∑(F_top50%_-F_BG_)/n). Our Matlab code provides a graphical user interface (GUI) to track the cell as it moves across the field of view, following a trajectory that is set by the waveform channel, to mark the location of 90° (maximal dorsal bend) or 270° (maximal ventral bend), and to trace ΔF/F against time or undulatory phase. We categorized traces from GABAergic motoneurons into low- and high-frequency (with 0.6 Hz as cut-off, see results) undulations of crawling inside the microfluidic channel; the undulation frequency was calculated as the fraction of locomotor cycle traveled by the cell of interest divided by the elapsed time. We collated and plotted all traces of ΔF/F against the phase of each type of cell at 5° bins. We corrected the soma position of motoneurons to their neuromuscular postsynaptic position according to the published perimotor locations (Chen *et al*., 2006; Haspel and O’Donovan, 2011). We removed videos or parts of videos in which the cell of interest went out of focus or if we could not positively identify it.

### Computational modeling

To probe different hypotheses about neural control and test their consequences in different fluid environments, we use an integrated neuromechanical model. Our model nematode consists of a smooth, continuous mechanical body, driven by neuromuscular activation, and subject to fluid drag forces (Denham *et al*., 2018). Our models aim to address the phenotypes of GABA transmission mutants during forward or backward locomotion, rather than directional changes or the *shrinker* phenotype. Specifically we focus on our results showing suppressed undulation frequencies in GABA mutants and suggesting that this phenotype is caused by defects in the ventral nerve cord circuitry, rather than by defects in premotor interneurons. To this end, we adapted an existing model of forward locomotion control (Denham *et al*., 2018). In particular, we explored two different paradigms of neural control: a proprioceptively driven model and a feedforward model, mimicking CPG control (Denham *et al*., 2018). We subjected each model to different perturbations and explored the behavior generated in our neuromechanical framework in virtual liquid and agar-like conditions. Model code and parameters are given in Code 8-1 and Table 8-1.

#### Neuromuscular model framework

Whether under CPG-, or proprioceptively-driven paradigms of locomotion control, our models of neural control generate a neural activation function that acts continuously along the body. We denote the neural activation by *A*(*u, t*), where *u* ∈ [0,1] represents the distance along the midline of the nematode from head (*u* = 0) to tail (*u* = 1), and *t* is time. This neural activation function corresponds to the difference between dorsal and ventral muscle inputs that arise from cholinergic B-type excitatory and GABAergic D-type inhibitory motoneurons on either side of the body. The mechanical component of the model is a flexible viscoelastic shell that is actuated by a spatially continuous muscle activation function, *β*(*u, t*), which is, in turn, derived from the neural activation function, *A*(*u, t*).

Our approach to modeling is parsimonious, with the aim of maximizing the explanatory power of the model with respect to the specific model hypotheses, and the effect of inhibition on undulation frequency. Therefore, asymmetries in dorsal versus ventral activation (Pierce-Shimomura, 2008), detailed muscle anatomy (e.g., around the pharynx and vulva), more detailed representations of neural connectivity (Haspel and O’Donovan, 2011), cuticle viscosity, muscle tone modulations, and rigorous models of the external fluid dynamics, are all neglected in this study. We also note that model results are qualitatively similar when using either a fixed or decaying amplitude of contractions along the body (see Boyle *et al*., 2012; Denham *et al*., 2018; Cohen and Ranner 2017). Accordingly, under both control paradigms, we consider uniform neural and muscle properties along the entire body.

#### Paradigm 1: CPG control

Following Denham *et al*. (2018) and in the absence of detailed CPG mechanisms, we model the essential feature of feedforward control, namely the existence of endogenous, central pattern generation that produces neural oscillations at a characteristic frequency and is embedded in a biomechanical body. We therefore model the neural activation by a retrograde traveling sine wave of unit amplitude with period, *T*_f_, and wavelength, *λ*_f_, that activates the muscles continuously along the body. Thus, the neural activation function takes the form

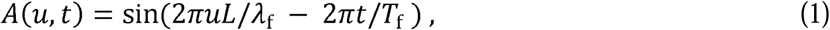

where *L* denotes the length of the body. While abstract, our CPG model is well suited to address the role of muscle inhibition, which has been and remains the main focus of research on the role of GABAergic motoneurons in *C. elegans* locomotion (McIntire *et al*., 1993; Schuske *et al*., 2004; Wu *et al*., 2018). As such, however, this model is agnostic about the role of neural inhibition in pattern generation itself.

#### Paradigm 2: Proprioceptive Control

In *C. elegans*, proprioceptive control of forward locomotion is mediated by B-type excitatory motoneurons (Wen *et al*., 2012; Fouad *et al*., 2018). Based on the neural circuitry, anatomy and ablation results (Chalfie *et al*., 1985; White *et al*., 1985; Chen *et al*., 2006) and consistent with previous models of forward locomotion (Niebur and Erdös, 1991; Cohen and Boyle 2010; Bryden and Cohen 2008; Boyle *et al*., 2012) in our model, B-type motoneurons receive a descending locomotion command input and local inhibitory input; internally, they are driven by a spatially extended proprioceptive input (Denham *et al*., 2018). For an in depth discussion of this class of models, see relevant reviews (Cohen and Boyle, 2010; Cohen and Sanders, 2014; Cohen and Denham, 2019).

#### Model B-type motoneurons

Following Boyle *et al*. (2012) and informed by electrophysiological recordings of bistable RMD head motoneurons (Mellem *et al*., 2008), we model excitatory ventral B-type (VB) and dorsal B-type (DB) motoneurons as bistable switches with binary states, {0,1}. Each neuron state is defined at every point along the body and denoted by *V*_VB_(*u, t*) and *V*_DB_(*u, t*). Neuronal bistability is implemented as distinct ON and OFF thresholds on the ventral (V) and dorsal (D) sides, 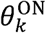 and 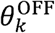 (*k* ∈ {V, D}), which need to be crossed for a neuronal state change of VB or DB to occur (akin to Boyle *et al*., 2012). Both VB and DB receive the same constant descending input from the AVB premotor interneurons, *I*_AVB_, which may be absorbed into the thresholds for parsimony. Proprioceptive inputs provide a feedback loop that couples the neural control and biomechanics. A proprioceptive input, *I*_*k*_(*u, t*), is defined for the dorsal side such that the input on the ventral side at the same position and time is −*I*_*k*_(*u, t*). Combined in our biomechanical framework, neuronal bistability and spatially extended proprioceptive inputs are sufficient to qualitatively capture observed gait modulation in response to changes in the external fluid environment, typically measured in terms of the frequency and waveform of undulations (Berri *et al*., 2009; Fang-Yen *et al*., 2010).

#### Model inhibition

As the limiting timescales of the dynamics are mechanical, we assume that D-type inhibitory motoneurons (with states *V*_VD_(*u, t*) and *V*_DD_(*u, t*)) respond instantaneously to excitation from B-type motoneurons on the opposite side of the body. Thus, if DB (or VB) is ON at some point along the body, VD (or DD) is activated. Following Boyle *et al*. (2012) and the published connectivity (Chen *et al*., 2006), we include VD-to-VB inhibition: Thus, VB is inhibited by VD (when DB is active), producing an input current *I*_VD_(*u, t*) = *w V*_VD_(*u, t*). In our implementation of the wild type model, we set the inhibition strength, *w*, such that the thresholds of VB and DB collapse to a single bistable switch when DB (hence also VD) is active. The corresponding reduced model, with only a single bistable switch, is presented and analyzed in depth in Denham *et al*., (2018).

Combining the above model components, VB and DB state switches are given by the following relations:

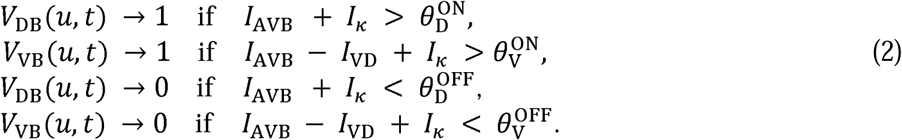

The state of a B-type neuron is driven by changes in proprioceptive input current, *I*_*k*_(*u, t*), which we define here as the mean of the body curvature, *k*(*u, t*), over a specified proprioceptive range, Δ(*u*), represented as a fraction of the body length:

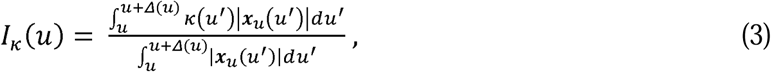

where *x* denotes a coordinate of a point along the body in the lab frame and the subscript, u, denotes a partial derivative and time dependence is implicit. This encoding of body posture is fed back to the motoneurons, thus closing the brain-body-environment loop. The above input current represents the effective stretch of the dorsal side of the body at that time. Here, we keep Δ (*u*) > 0 which by our convention corresponds to a strictly posterior proprioceptive range, though Δ (*u*) < 0, and combinations of anterior and posterior range are also possible (see Denham *et al*., 2018). For *u* sufficiently close to the tail, the proprioceptive range terminates at the tail, therefore decreasing linearly with *u*. Hence the denominator in Equation (3) normalizes by this range, effectively making posterior neurons more sensitive to stretch ‘density’ to compensate for the reduced range. In all simulations, we set the proprioceptive range to half the body length, posteriorly to the muscle coordinate (Boyle *et al*., 2012; Denham *et al*., 2018).

As our biomechanical model uses a midline parametrization to determine the internal forces driving body curvature, our neural activation function sums the contributions of local neurons from both sides of the body, weighted by excitatory and inhibitory neuromuscular junction weights, 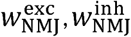, according to

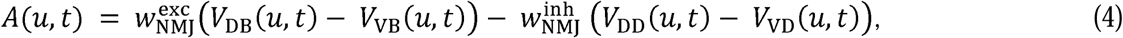

to define our activation, *A*(*u, t*), which then drives the muscle via Equation (5) below.

#### Model muscles

Under proprioceptive control, body wall muscles are modeled by a leaky integration equation that converts the neural activation function at every point along the body, *A*(*u, t*), to a mechanical torque. Specifically, muscles receive local inputs from ventral and dorsal motoneurons and output the muscle activation, *β*(*u, t*), acting on the animal’s midline. The muscle model contains two free parameters: a muscle time scale, *τ*_m_, set to 100 ms (Table 8-1) in the wild type, and an amplitude or maximum curvature (10 mm^-1^ in the wild type, see Denham *et al*. (2018) for discussion). Thus, given the neural activation, *A*(*u, t*), the muscle response is given by

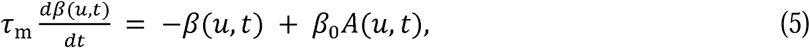

where the muscle activation, *β* (*u, t*), corresponds to a preferred curvature (akin to a rest length of a spring) and *β*_0_ sets the curvature amplitude (Equation (8) below).

In the case of CPG control, and in order to accommodate the disinhibition of body wall muscles, we assume that the muscle activation in the wild type depolarizes more sharply than in GABA mutants. We therefore set the muscle activation to

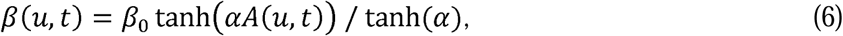

with the steepness, *α*, corresponding to the strength of the muscle disinhibition and where the denominator normalizes the amplitude of the muscle activation.

#### Biomechanics

Our model of the *C. elegans* body consists of a thin incompressible viscoelastic shell, subject to internal pressure and external forces (see Cohen and Ranner (2017) and Ranner (2020) for a detailed description of the body mechanics, mathematical derivation and numerical methods). At each point along the body, we assume the radius of the nematode’s body is fixed in time, which allows us to collapse all internal and external forces onto the midline (Denham *et al*., 2018; Ranner, 2020). Four free parameters modulate the biomechanical properties of the body and its interaction with the environment. The body is parametrized by its effective Young’s modulus, *E* (the effective elastic resistance to bending or body stiffness, set to 100 kPa in the wild type), and an internal viscosity (the body damping in response to bending), which we neglect for simplicity. Environmental forces are modeled by resistive force theory and parametrized by tangential and normal drag coefficients, *K*_*τ*_ and *K*_*v*_, that act against the cuticle to resist the motion in the respective directions, denoted by unit vectors, ***τ*** and ***v*** (Berri *et al*., 2009; Boyle *et al*., 2012; Denham *et al*., 2018). Accordingly, the resistive forces are given by

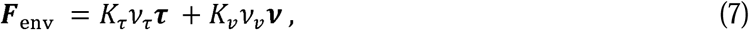

where *v*_*τ*_ and *v*_*V*_ represent the normal and tangential components of the velocity of a point on the surface of the body. This allows us to model both Newtonian and linear viscoelastic environments in the low Reynolds number regime. Our simulations here mimic two environmental conditions: swimming in liquid (buffer solution) and crawling on agar. The linear viscoelastic approximation of the fluid dynamics appears valid for these purposes (Berri *et al*., 2009), provides computational efficiency and is sufficient to reproduce the experimentally observed gait transition under models of feedback control (Boyle *et al*., 2012; Denham *et al*., 2018).

The model equations balance internal and external forces and torques subject to mass conservation within the nematode’s body and an inextensibility closing constraint (Cohen and Ranner, 2017; Ranner, 2020). To produce a body bend, the signed midline curvature, *k*(*u, t*), is driven by the muscle activation, *β* (*u, t*), (here, a torque that is represented as a preferred body curvature) which in turn is driven by neural activation, *A*(*u, t*), in either the CPG or proprioceptive control paradigm. By convention, positive and negative *β* (*u, t*) and *k*(*u, t*) correspond to dorsal and ventral excitation and body curvature, respectively. The timescale of undulations depends on both the elasticity of the body, E, and the resistivity of the environment (Cohen and Ranner, 2017; Denham *et al*., 2018). The balance of forces is summarized as follows:

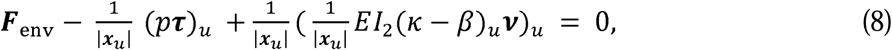

where *p* represents the nematode’s internal pressure which acts as a line tension along the midline of the nematode and *I*_2_ denotes the second moment of area. Zero force and zero torque are enforced at the boundaries, such that *β*(*u, t*) = *k*(*u, t*) at both the head and tail ends of the body (*u* = 0,1). We note that our biomechanical model is symmetrical across the dorsal and ventral sides.

#### Model perturbations

Based on our experimental results, we explore three hypotheses to explain the role of inhibition in sustaining high frequency locomotion; each of these hypotheses are tested separately in our computational models under purely feedforward (CPG) and purely feedback (proprioceptive) neural control. To qualitatively describe different perturbations, relatively small perturbations are the most useful in that they are restricted to the domain in which the model was set (best matching crawling and swimming behaviors of the wild type). To determine the range of perturbations and their effects, parameter sweeps were performed for hypotheses 1 and 2 (Ext. Fig. 8-1). Unless otherwise stated, we only knock out one form of inhibition in the model in a given simulation.

### Hypothesis 1

Cross-inhibition of the opposing body wall muscles increases the dorsoventral difference in muscle activation. To implement this hypothesis in our model, we eliminated all muscle inhibition by D-type neurons, leaving neural inhibition intact. In our model, the muscle forcing on both sides of the body is combined into a single muscle activation function acting along the midline of the body. Therefore, the elimination of muscle cross-inhibition by D-type neurons effectively results in reduced activation on the bending side. In practice, we implement this change with a multiplicative constant applied to our preferred curvature:

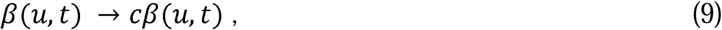

with *c* = 1 in the wild type and *c* = 0.8 in the default perturbation. In the absence of cross muscle inhibition, setting *c* < 1 indicates that the muscle on the stretched side remains partially activated (Goodman *et al*. 2011; Boyle *et al*. 2012). In our proprioceptively driven model, we set the neuromuscular junctions of B- and D-type motoneurons such that eliminating muscle inhibition gives an identical outcome (see Table 8-1).

### Hypothesis 2

Disinhibition of body wall muscles should increase muscle force, immediately after muscle inhibition is released, resulting in a sharper upstroke in muscle activation on the contracting (bending) side of the body. We distinguish between CPG and proprioceptively driven control as follows:

#### CPG control

In the wild-type model, we enforce a sharpness in the upstroke of muscle drive using a sigmoidal muscle activation function, which we neglect in disinhibition deficient models:

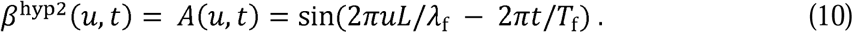

More generally, for the purpose of parameter sweeps (Ext. Fig. 8-1), we vary the steepness of the upstroke, *α*, in Equation (6).

#### Proprioceptive control

While this could be modeled by a more complex nonlinear muscle model, we reasoned that the effect of a muscle disinhibition can be qualitatively modeled using our linear muscle equation by allowing disinhibition deficient models to have a slower muscle timescale on the contracting side of the body. Due to the alternation of muscle contraction, the combined dorsoventral muscle activation, *β*^hyp2^(*u, t*), of the midline will experience an overall slower effective muscle timescale, 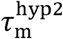, yielding

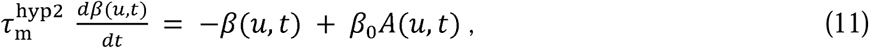

where 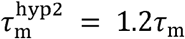.

### Hypothesis 3

Inhibitory reset of VA and VB by VD is required for rapid swimming. Boyle *et al*. (2012) hypothesized that nematodes lacking neural inhibition will fail to undulate in water-like environments, in which wild-type nematodes undulate at higher frequency. As this hypothesis depends explicitly on the GABAergic neural circuit (and as our simplistic feedforward model does not take into account individual motoneurons), we only explore this hypothesis in our proprioceptively driven model. In this model, VB activation depends on the state of DB due to the DB→VD →VB connectivity. Elimination of neural inhibition implies that VB neurons are not inhibited when DB neurons are active, but that muscle cross-inhibition remains intact. To model defects corresponding to this hypothesis, we set *I*_VD_ = 0 at all times. This effectively results in a constant positive offset applied to both thresholds of the VB bistable switch.

#### Parameter choices for perturbations

As the change of waveform in GABA mutants is subtle, and extensive literature points only to subtle changes in forward locomotion (McIntyre *et al*., 1993; Schuske *et al*., 2004; Bamber *et al*., 2005, and our own data herein), we assume that the effect of excitation is much stronger than that of inhibition. Muscle contraction typically leads to, at most, a 20% change in muscle length (Boyle *et al*., 2012; Petzold *et al*., 2011). Therefore, we set 20% as an upper cap on perturbation strength in all model perturbations. We note that neural activation in our model maps linearly to muscle input and muscle length. Applying the same magnitude perturbations in all cases aids intuition and points to the robustness of the mechanisms investigated in the model. Thus, our choice of 20% is given as a visual example in the figures for a particular parameter value and does not constitute a quantitative model prediction. Values up to 20% gave qualitatively similar results (Ext. Fig. 8-1).

##### Sample size, experimental design, and statistical analysis

For free crawling behavior tracking of N2 (wild type), the number of videos, animals, and data points were 12, 235, and 67,209, respectively, for VC1433, the numbers were 11, 213, and 80,407; for TM5916 the numbers were 11, 255, and 94,446, and for TM5487 the numbers were 12, 289, and 121,374. For free swimming behavior tracking of N2 (wild type), the numbers of videos, animals, and data points were 46, 707, and 164.057, respectively, for VC1433, the numbers were 39, 778, and 247,656; for TM5916 the numbers were 38, 1014, and 316,094; and for TM5487, the numbers were 34, 668, and 293,482. For free crawling behavior tracking of TOL12 without ATR feeding, the number of videos, animals, and data points were 20, 375, and 104,098, respectively. For TOL12 that were fed with ATR and tracked in the dark, the numbers were 27, 335, and 98,017; and for TOL12 that were fed with ATR and were exposed to lime light during tracking, the numbers were 26, 235, and 66,803. For free swimming behavior tracking of TOL12 without feeding ATR, the number of videos, animals, and data points were 20, 326, and 81,291, respectively. For TOL12 that were fed with ATR in advance and tracked in the dark the numbers were 22, 361, and 97,630; and for TOL12 that were fed with ATR and were exposed to lime light during tracking, the numbers were 18, 216, and 48,323. We collated the means of these parameters from individual animals from all the videos, and then used one-way ANOVA and Tukey’s multiple comparison to compare the estimated population means among strains or different conditions.

For the head and tail harsh touch experiments, we obtained 9 trimmed videos for each stimulation direction for N2 and GABA transmission knockout strains (VC1433, TM5916, TM5487) as well as 9 trimmed videos for *vab-7* mutants (CB1562) of head harsh touch and *unc-4* mutants (CB120) of tail harsh touch. We used a paired two-tail t-test to determine statistical differences in the parameters before and after stimulation of the same strain, and one-way ANOVA and Tukey’s multiple comparison for the parameters of after harsh touch to the head or tail between wild-type and mutant strains, including GABA transmission knockouts, and *vab-7* and *unc-4* mutants.

To generate linear regressions of sinusoidal crawling and swimming behaviors, we first picked the data points where the primary or secondary wavelengths could be extracted by the analysis software (Tierpsy). These data points in crawling behavior comprised 4% to 23% of the raw data. The sample sizes of wild type, *unc-25, unc-46*, and *unc-49* during forward crawling were 11,935 (23%), 7,760 (14%), 7,773 (12%), and 5,378 (7%), respectively; those during backward crawling were 1,260 (11%), 2,319 (11%), 2,552 (10%), and 1,714 (4%), respectively. The data points in swimming behavior comprised 6% to 54% of the raw data. The sample sizes of wild type, *unc-25, unc-46*, and *unc-49* during forward swimming were 74,948 (54%), 46,355 (26%), 69,682 (31%), and 17,015 (11%), respectively; those during backward swimming were 2,137 (14%), 6,249 (11%), 16,728 (21%), and 7,297 (6%), respectively. Then, we picked the data points where the path of an undulation cycle (translocation speed × undulation cycle period) was larger than either the primary or secondary wavelength. Based on this selection criterion, we picked 27-54% data points for the correlation construction in crawling behavior. The sample sizes of wild type, *unc-25, unc-46*, and *unc-49* during forward crawling comprised 5,671 (48%), 2,058 (27%), 2,259 (30%), and 2,313 (43%), respectively; those during backward crawling were 686 (54%), 1,096 (47%), 1,199 (47%), and 856 (50%), respectively. We picked 0.4% to 26% data points in swimming behavior. The sample sizes of wild type, *unc-25, unc-46*, and *unc-49* during forward swimming were 271 (0.4%), 3,985 (14%), 2,086 (3%), and 2,965 (17%), respectively; those during backward swimming were 138 (7%), 1,150 (18%), 975 (6%), and 1,897 (3%), respectively. Using these selected data, we calculated linear regressions between undulation frequency and translocation speed for each of the free locomotion conditions, forward and backward crawling and swimming separately.

For the calcium imaging experiment, we included data analyzed from 11 focused and cell-identifiable videos of TOL15; 11 and 9 videos of CZ1632, TOL11 in 0.5% (w/w) methyl cellulose, respectively, and 12 and 9 videos in 3% (w/w) methyl cellulose, respectively. From these videos, we analyzed 27 and 24 ventral and dorsal body-wall muscles, respectively, in TOL15; 17 VD and 18 DD motoneurons in the high undulation frequency and 26 VD and 13 DD motoneurons in low frequency in CZ1632; and 32 VD and 15 DD motoneurons in the high undulation frequency and 27 VD and 14 DD motoneurons in low frequency in TOL11.

##### Code Accessibility

The code described in the paper is freely available online at [URL redacted for double-blind review]. The code is available as Extended Data.

## Results

### Both wild-type and GABA transmission mutants shrank in response to harsh touch to the head or tail

*C. elegans* mutants defective in GABA transmission are known as ‘*shrinkers*’ because of their distinct decrease in body length in response to a harsh touch to the head (McIntire *et al*., 1993a; McIntire *et al*., 1993b; Schuske *et al*., 2004), to which wild-type animals respond with rapid backward motion. Because this phenotype has been described as a defect in backward locomotion, we first looked at the shrinking response of wild-type and GABA transmission mutant animals to harsh touch to the head, which induces rapid backward undulation in wild-type animals as well as the response to the tail touch to study the association of shrinking response to forward locomotion. We chose three knockout mutant strains in which hundreds to thousands of DNA base pairs had been deleted in the genes encoding the GABA synthesis enzyme glutamic acid decarboxylase (GAD, *unc-25*, VC1433), vesicular transporter (*unc-46*, TM5916), and ionotropic GABA receptor (*unc-49*, TM5487), and compared them to the laboratory reference strain (wild type, N2). We found that the shrinking response was common in wild-type and mutant animals following head or tail stimulation (Fig. 1a). To quantify the shrinking response of wild-type and the GABA transmission knockout animals, we measured the change in body length for each strain and stimulation direction (n=9 each) at four different times (before, immediately after, 0.5 s, and 2 s after stimuli), and normalized the body length to their value before stimuli (Fig. 2a, Table 2-1). Immediately after a harsh touch to the head or tail, both wild-type and the GABA transmission knockout animals decreased their body length: the body length of wild-type animals reduced to 94 ± 2% and 95 ± 3% (mean ± SD), respectively; while the body length of GABA transmission knockouts was reduced to 90-91 ± 2-4% and 91-92 ± 3-4% immediately after harsh touch to the head or tail, respectively across the different strains. In addition, wild-type animals recovered to their pre-stimulus body length faster. At 2 s after the head or tail stimulation, the body length of wild-type animals reached 98 ± 4% and 99 ± 2% of the pre-stimulus body length; while that of GABA transmission knockouts reached 92-94 ± 4-6% and 93-97 ± 2-5%, respectively across the different strains (Fig. 2a, Table 2-1). Moreover, GABA transmission knockouts occasionally exhibited spontaneous shrinking (Fig. 1f). The brief shrinking of wild-type animals was only associated with vigorous escape response from strong aversive stimuli such as harsh touch and a strong chemical repellent (97 ± 1%, 50 mM CuSO_4_, on-food; Table 2-1), but not with a weak chemical repellent (99 ± 1%, 0.1% SDS, off-food; Table 2-1).

**Figure 1.**
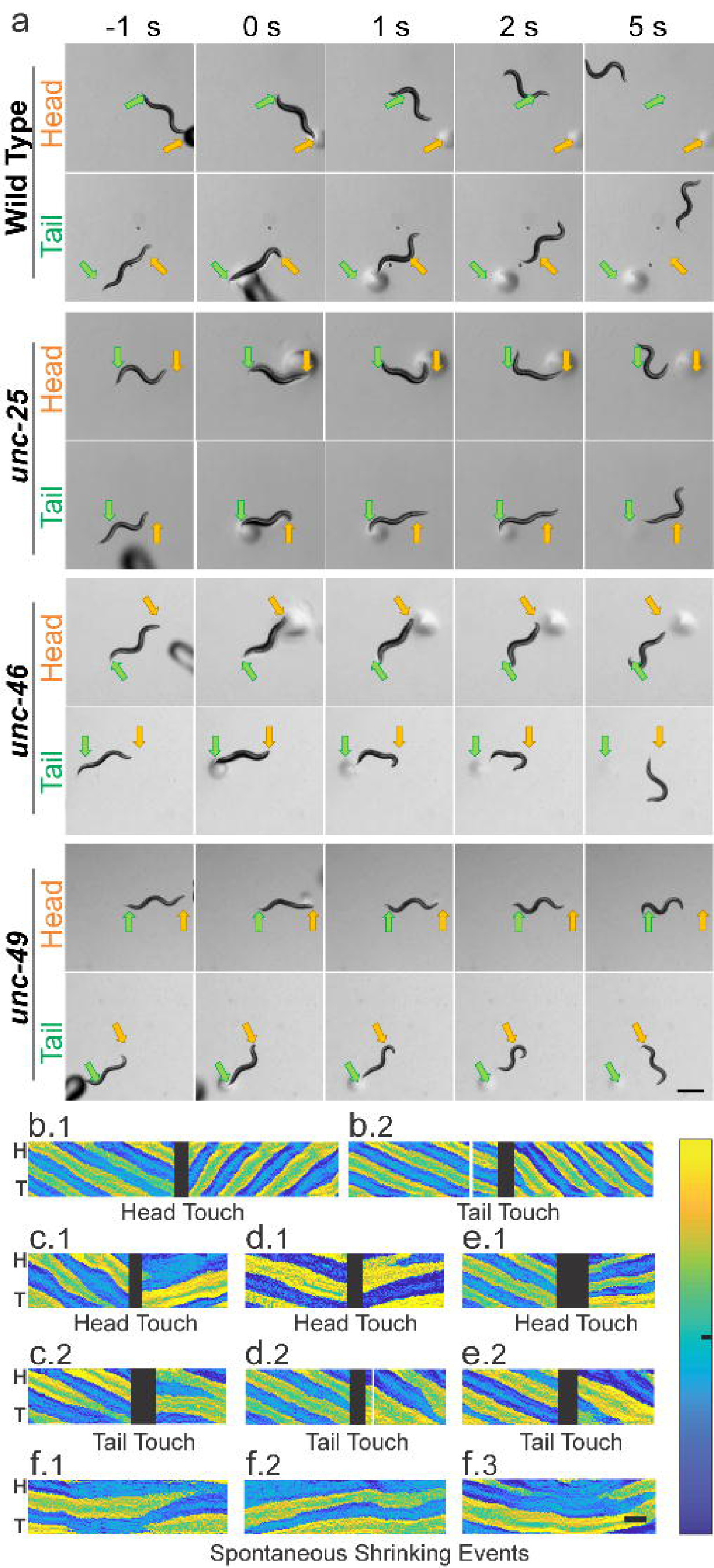
Wild type and GABA transmission knockouts shrank after harsh touch to the head or tail, while wild-type animals escaped rapidly. (a) While moving forward, animals were touched by a blunt glass probe as a harsh stimulation to the head (yellow arrow) or tail (green arrow). Wild-type animals briefly shrank immediately after stimulation (compare 0 s to -1 s before stimulation) and rapidly moved away from the stimulus, while GABA transmission knockouts shrank for a longer time and moved away slowly. Note that by 5 s all animals had moved away from the stimuli. Common scale bar is 0.5 mm. Dark and circular shadows are the glass probe and the marks it left on agar. After harsh stimulation, wild-type animals (b) increased undulation frequency to escape backwards (b.1) or forwards (b.2). In contrast, all three GABA transmission mutants – *unc-25* (c), *unc-46* (d) and *unc-49* (e) – decreased undulation frequency when they moved away from the stimuli. Moreover, there was a period in which the body posture of these mutants did not change after the stimulation (c-e). (f) The mutants, *unc-25* (f.1), *unc-46* (f.2) and *unc-49* (f.3), occasionally shrank without stimulation. Yellow and blue shaded areas in b-f represent dorsoventral curvature (vertical axis is the length of animal from the head, H, on the top to the tail, T, color bar is curvature -10 to 10 mm^-1^, indifferent to dorsoventrality), along time (horizontal axis, common scale bar is 1 s); black blocks indicate gap in tracking during the touch stimulation.

**Figure 2.**
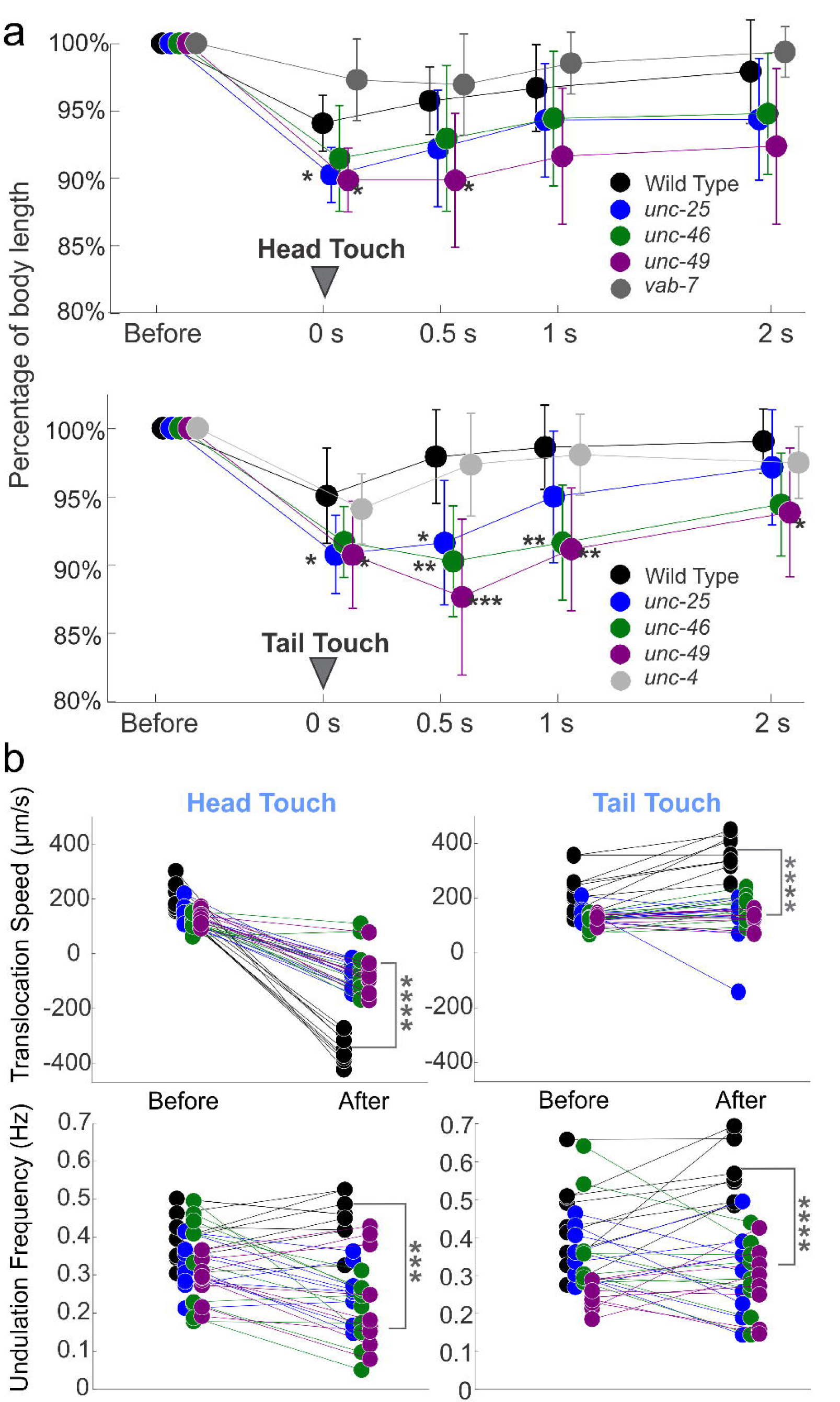
Both wild type and GABA transmission knockout animals shrank after harsh touch stimuli; wild-type animals shrank less, recovered sooner, and crawled away more rapidly than knockout animals. (a) Wild type and GABA transmission knockout animals decreased their body length after harsh stimuli to the head or tail (0 s). Compared to wild type, GABA transmission mutants decreased their body length more and recovered more slowly (0.5, 1, and 2 s after stimuli). The body length change of *vab-7* and *unc-4* were not statistically different from that of wild type. Body length is normalized to pre-stimulus value for each animal. See Table 2-1 for details. (b) Wild-type animals moved with higher mean undulation frequency and mean translocation speed during 5 s post-stimulus compared to the 5 s pre-stimulus, while the mutants did not. Two *unc-46* mutant animals did not change locomotion direction after harsh head touch, neither did one *unc-25* mutant animal after harsh tail touch. One-way ANOVA with Tukey’s pairwise comparison to wild type, * p <0.05, ** p<0.01, *** p<0.001, **** p<0.0001. Black circles: wild type, blue: *unc-25*, green: *unc-46*, and purple: *unc-49*. See Tables 2-2 and 2-3 for details.

A possible explanation for the dorsoventral coactivation of body-wall musculature that causes body shrinkage in response to harsh touch is the simultaneous activation of all A- and B-type cholinergic motoneurons by premotor interneurons that are associated with forward and backward locomotion. These distinct groups of interneurons activate either, DA and VA motoneurons during backward, or DB and VB motoneurons during forward locomotion (Haspel *et al*., 2010). Thus, we hypothesize that shrinking results from a short temporal overlap of activity of A- and B-type cholinergic motoneurons. To test this hypothesis, we took advantage of developmental phenotypes that render mutants in a complementary pair of homeodomain transcription factors (UNC-4 and VAB-7), each lacking either A- or B-type motoneurons. In *unc-4* mutant animals (CB120), cells destined to be A-type motoneurons differentiate as B-type motoneurons and receive input from forward-associated premotor interneuron (Miller *et al*., 1992; Miller and Niemeyer, 1995). While it is unclear whether the misdifferentiated neurons fully function as B-type motoneurons, their function as A-type is impaired. Consequently, only their backward locomotion in uncoordinated. Hence, we tested whether *unc-4* mutant animals shrink in response to harsh touch to the tail that induces an acceleration forward, towards their unaffected direction. Correspondingly, in *vab-7* mutant animals (CB1562), cells destined to be DB motoneurons differentiate as DA motoneurons (Esmaeili *et al*., 2002). Consequently, only their forward locomotion is uncoordinated. Hence, we tested whether *vab-7* mutant animals shrink in response to harsh touch to the head that induces an acceleration towards their unaffected direction. The harsh touch stimuli induced a brief reduction of body length in animals of both mutant strains (Fig. 2a) that was not different from that of the wild type (Table 2-1). These results are inconsistent with the hypothesis of coactivation by premotor interneurons, suggesting instead that the shrinking phenotype is localized within the VNC motor circuit and not associated with directional change.

### GABA transmission knockouts failed to escape as rapidly as wild type after harsh touch stimulation

After harsh touch stimulation, wild-type animals were able to escape at high frequency and translocation speed, while the mutants moved away slowly (Fig. 1a, note location at 5 s). We measured the translocation speed and midbody undulation frequency as well as the maximal amplitude and primary wavelength, within 5 s before and 5 s after stimulation (n=9 video clips for each strain and stimulation location, Fig. 2b, Tables 2-2 and 2-3). After harsh touch to the head or tail, wild-type animals escaped with significantly higher translocation speed and undulation frequency, while GABA transmission knockouts had lower mean translocation speeds except for *unc-46* and *unc-49* after harsh tail touch, and lower mean undulation frequency, except for *unc-49* after harsh tail touch. Compared to the locomotion within 5 s after the harsh head or tail touch of the wild type, all the GABA transmission knockouts had significantly lower translocation speed (one-way ANOVA with Tukey test, p <0.0001 pairwise comparison to wild type) and undulation frequency (p <0.05 pairwise comparison to wild type; Fig. 2b, Table 2-3). Wild-type and GABA transmission knockouts exhibited larger amplitudes, but unchanged wavelengths after stimulation statistically (except for *unc-46* after tail harsh touch, Fig. 2b, Table 2-2). Note that to quantify the shrinking response, we applied harsh touch to sinusoidally forward crawling animals (both wild type and mutants) and discarded assays that did not stand to this criterion. A possible bias from this selection underestimates the actual differences among strains.

### Defect in GABA transmission resulted in slow locomotion during free crawling and swimming

Early descriptions of the locomotion defect of GABA transmission mutants and animals in which most GABAergic motoneurons have been laser-ablated emphasize the absence of backward locomotion (McIntire *et al*., 1993b; Yanik *et al*., 2004). However, we noted that GABA transmission knockout strains seem to crawl at low undulation frequencies in both directions, and their locomotion was more impaired during swimming in liquid (consistent with Boyle *et al*. (2012)). Moreover, shrinking is not unique to backward locomotion but occurs in both mutants and wild-type animals regardless of direction. We recorded and compared unstimulated free crawling and swimming behaviors of wild-type and GABA transmission knockout strains as well as an optogenetic strain (TOL12) in which GABAergic motoneurons can be inactivated acutely by light.

Crawling on the NGM agar surface without bacterial lawn, GABA transmission knockout strains crawled slowly. The distribution of translocation speeds and frequencies of GABA transmission knockouts during forward and backward crawling were significantly different from those of the wild type (KS test, p <0.0001, for *unc-25, unc-46*, and *unc-49*). The wild-type distribution of undulation frequency was bimodal for both forward and backward crawling. The undulation frequency during forward crawling fits two Gaussian distributions: a rapid distribution with 0.84 mixing proportion has a mean value of 0.47 Hz, and a slow one with 0.16 mixing proportion has a mean value of 0.17 Hz (Fig. 3a). GABA transmission knockout strains crawled at lower mean translocation speed and mean undulation frequency compared to the wild type (Fig. 3, Table 3-1). Only the mean undulation frequency of VC1433 during forward crawling was not significantly lower than that of the wild type. The maximal amplitudes and primary wavelength of GABA transmission knockouts were significantly smaller than those of the wild type.

**Figure 3.**
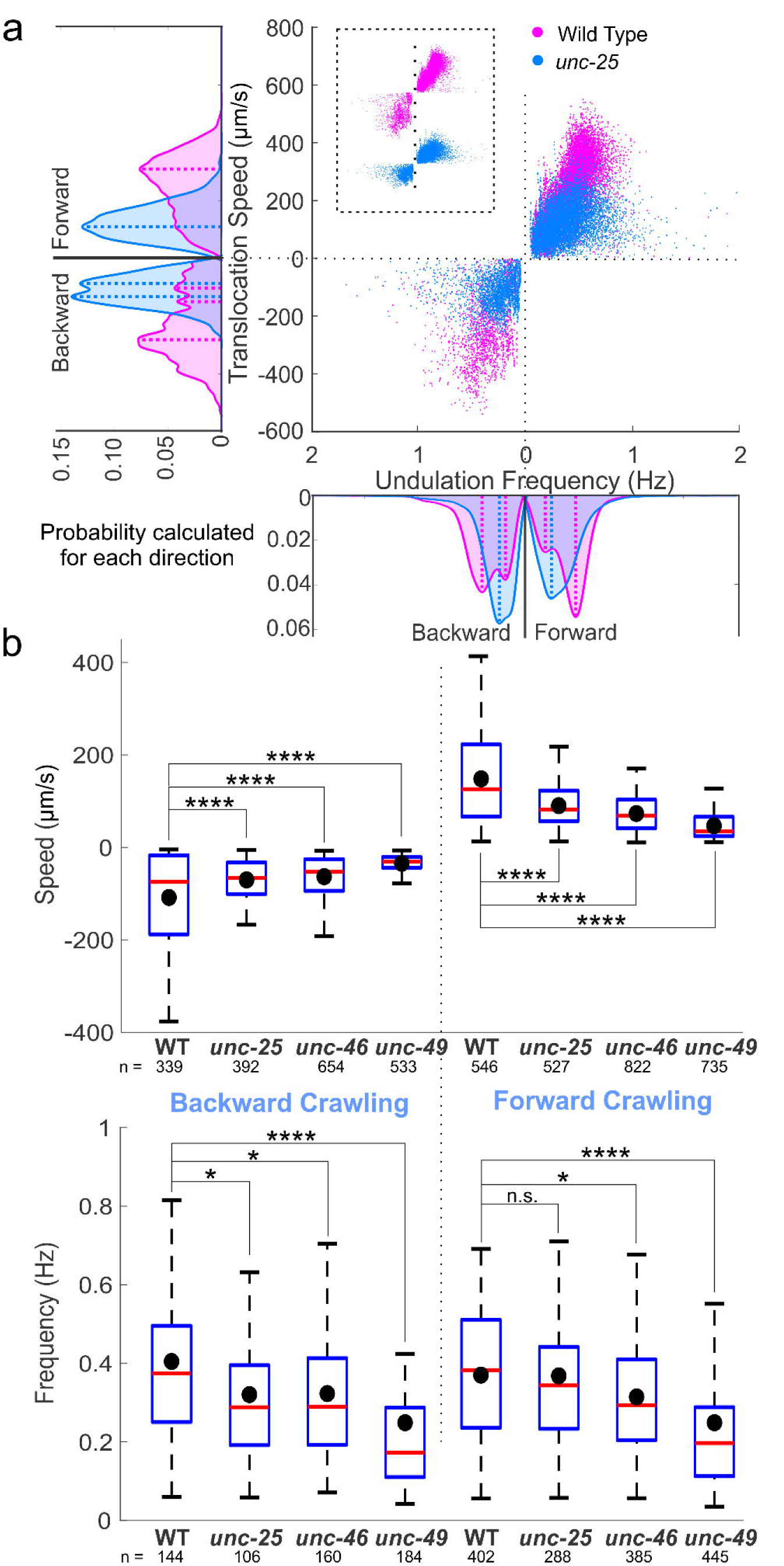
GABA transmission mutants crawled more slowly than wild type. (a) *unc-25* knockout animals crawled on agar surface with lower translocation speed and undulation frequency than wild type. Negative translocation speed and undulation frequency left (and right) of 0 represent backward (and forward) locomotion. Area plots show the probability of occurrence for each direction with dashed lines indicating the main peaks in the area plots. Probabilities were generated from histograms with bin size of 20 µm/s and 0.02 Hz translocation speed and undulation frequency, respectively. (b) Translocation speeds of all the GABA transmission knockouts were significantly lower than that of wild type (WT) during forward and backward crawling. The undulation frequencies of GABA transmission knockouts were significantly lower than those of wild-type animals, except for that of *unc-25* during forward crawling. Blue boxes: first to third quartiles of each dataset; red lines: medians; black dots: mean values; whiskers: minimum and maximum, excluding outliers (beyond 1.5-fold interquartile range from middle 50% data). ANOVA with Tukey pairwise comparison to wild type, n.s. p>0.05, * p<0.05, *** p<0.001, **** p<0.0001. See Table 3-1 for details.

During swimming, *C. elegans* undulates at higher frequencies, wavelengths, and amplitudes (Karbowski *et al*., 2006; Pierce-Shimomura *et al*., 2008; Butler *et al*., 2015; Berri *et al*., 2009, Fang-Yen *et al*., 2010). If the absence of GABA transmission impairs fast undulations, it should have a more severe effect on swimming behavior with a more distinct difference between mutants and wild type. We therefore recorded the swimming behavior of wild-type and GABA transmission knockouts in NGM buffer. Wild-type animals alternated dorsal and ventral bends evenly with larger undulation amplitude and wavelength than during crawling (Fig. 4a1). In comparison, GABA transmission knockouts undulated at a lower frequency with dorsoventral body alternation that was not as symmetric as that of the wild type (Fig. 4a2). The distribution of translocation speeds and frequencies of GABA transmission knockouts during forward and backward swimming were significantly different from those of the wild type (KS test, p<0.001; for *unc-25, unc-46*, and *unc-49*; Fig. 4b).

**Figure 4.**
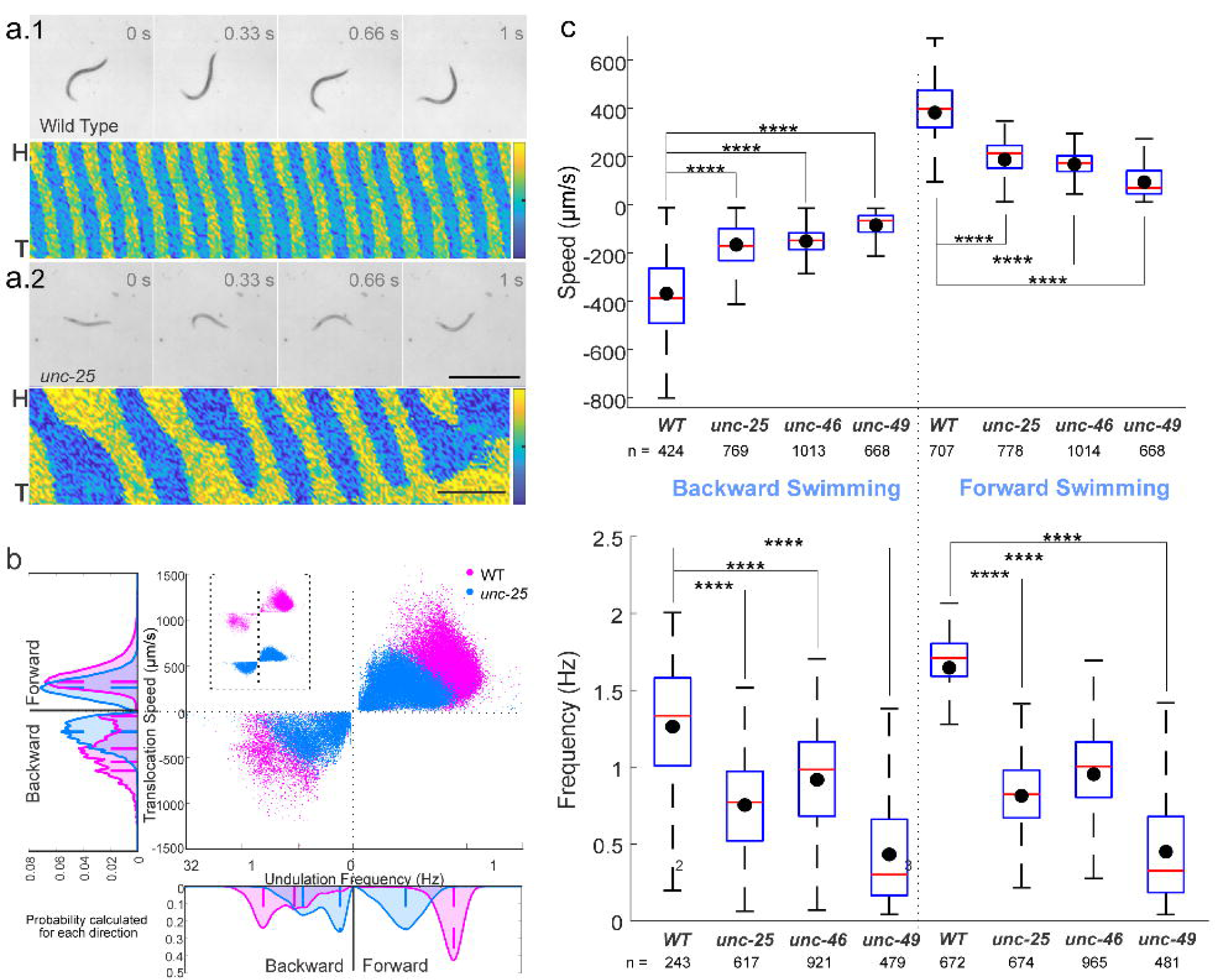
GABA transmission knockouts swam more slowly than wild type. Compared to wild type (a.1), *unc-25* knockout (a.2) in liquid swam with lower undulation frequency and translocation speed, as well as smaller undulation amplitude and wavelength. Video frames are 0.33 s apart, common scale bar is 1 mm; yellow and blue shaded areas of kymograms represent dorsoventral curvature (vertical axis is the length of animal from the head, H, on the top to the tail, T, color bar is curvature -10 to 10 mm^-1^, indifferent to dorsoventrality), along time (horizontal axis, scale bar is 1 s). (b) *unc-25* knockout animals swam slower with lower translocation speed and undulation frequency than wild type. Area plots show the probability of occurrence for each direction with dashed lines indicating the main peaks in the area plots. Probabilities were generated from histograms with bin size of 24 µm/s and 0.05 Hz translation speed and undulation frequency, respectively. (c) Translocation speeds and undulation frequencies of all the GABA transmission knockouts were significantly lower than those of wild type (WT) during forward and backward swimming. Box plots as in Fig. 3. ANOVA with Tukey pairwise comparison to wild type, **** p<0.0001. See Table 4-1 for details.

Indeed, GABA transmission knockout strains swam at a significantly lower translocation speed and undulation frequency than the wild type (Fig. 4bc, Table 3-1). The maximal amplitudes and primary wavelength of GABA transmission knockouts were significantly smaller than those of the wild type.

The GABA transmission knockout animals are chronically impaired, and the effects we describe on locomotion could rise during development or due to compensation mechanisms. Therefore, we studied the locomotion of a transgenic strain (TOL12) that expresses Archaerhodopsin-3 (Arch3) on the cellular membrane of GABAergic motoneurons, so they can be acutely inactivated by light. Arch3 is a light-sensitive proton pump that requires the cofactor all-trans-retinal (ATR) and causes hyperpolarization when exposed to lime-color light (Nagel *et al*., 2005; Okazaki *et al*., 2012). We fed transgenic animals with ATR for 24 h and tracked their swimming or crawling behaviors under infrared light (that does not activate Arch3) and then under lime-color light, which is the optimal activation wavelength of Arch3 (Mattis *et al*., 2011; Okazaki *et al*., 2012). To prevent desensitization of Arch3 after long exposure, we limited lime-color light to 1 minute (Okazaki *et al*., 2012). We also tracked the locomotion behavior of transgenic animals that were not fed ATR as a negative control. When GABA transmission was inactivated acutely by hyperpolarizing GABAergic motoneurons, the frequency of undulations decreased, comparable to the effect of GABA transmission mutants. Swimming under lime-color light, the transgenic animals fed with ATR were slower and less coordinated, and their maximal amplitude and primary wavelength decreased, but quickly recovered when the optogenetic activation light was turned off (Fig. 5abc). Crawling on agar surface under lime-color light, the translocation speed of transgenic animals fed with ATR was significantly lower than that of the same animals under infrared light and that of the animals not fed ATR under lime-color light. Their undulation frequency was lower than that of the same animals under infrared light and that of the animals not fed ATR. Swimming in NGM buffer, their translocation speed was lower (p < 0.0001) than that of the same animals under infrared light and that of the animals not fed ATR. Their undulation frequency was lower (p < 0.0001) than that of the same animals under infrared light and that of the animals not fed ATR (Fig. 5d, Table 5-1). In contrast with the swimming of GABA transmission mutants, some data points are as high frequency and fast as in control animals (compare Fig. 4b to Fig. 5c), suggesting an incomplete optogenetic neuronal inactivation.

**Figure 5.**
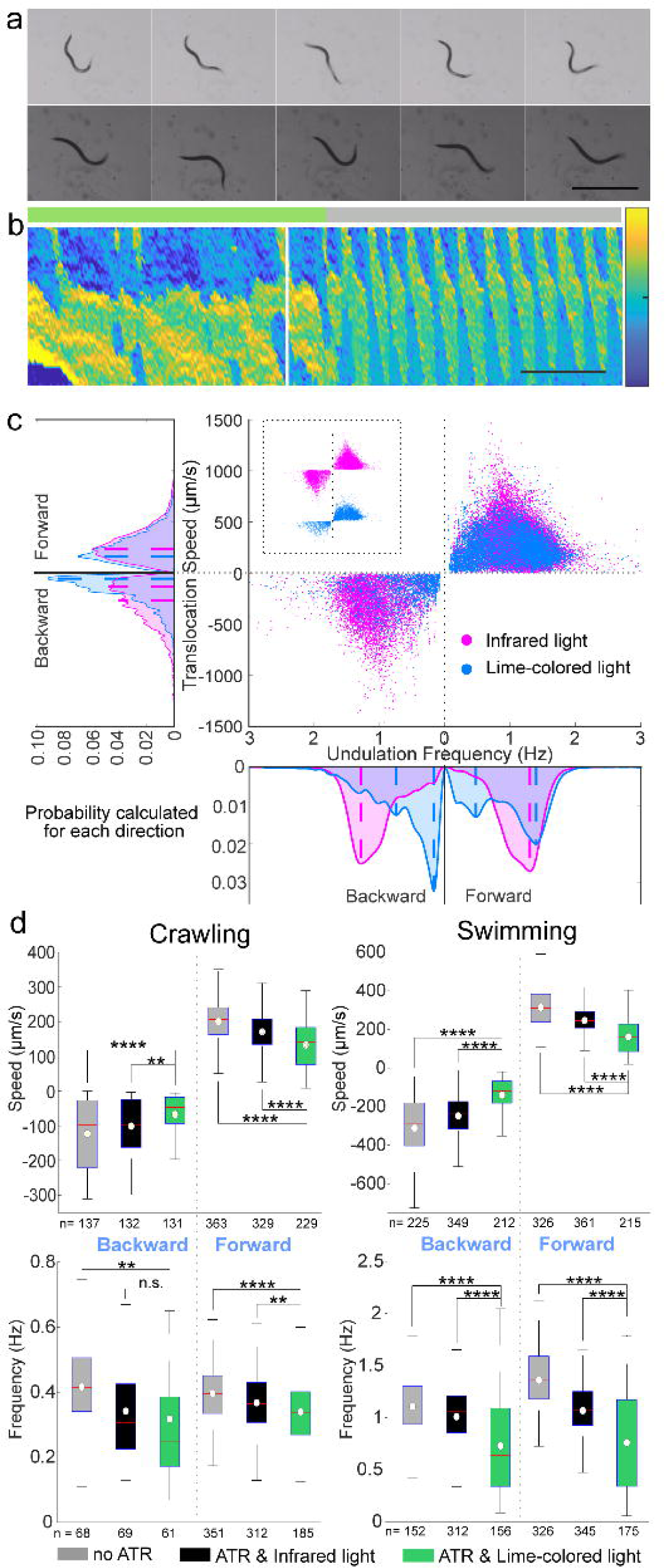
Acute optogenetic inactivation in GABAergic motoneurons induced slow locomotion during free swimming and crawling. (a) When GABAergic motoneurons were inactivated optogenetically (top framed and green line on top of kymogram) while swimming in liquid, undulations were slower and more uncoordinated. When the optogenetic inhibition was withdrawn (bottom frames and grey line on top of kymogram), the animal regained coordinated undulatory swimming and its undulation frequency doubled from 0.5 to 1 Hz. Postures shown at 0.33 s intervals; scale bar is 1 mm; yellow and blue shaded areas of kymograms represent dorsoventral curvature (vertical axis is the length of animal from the head on the top to the tail, color bar is curvature -10 to 10 mm^-1^, indifferent to dorsoventrality), along time (horizontal axis, scale bar is 1 s). (b) The optogenetic strain fed with ATR swam slower when optogenetically inactivated (by lime-colored light) compared to the same animals under infrared light. Area plots show the probability of occurrence for each direction with dashed lines indicating the main peaks in the area plots. Probabilities were generated from histograms with bin size of 24 µm/s and 0.05 Hz translation speed and undulation frequency, respectively. (c) When GABAergic motoneurons were acutely inhibited (green boxes, ATR and lime-colored light), animals crawled and swam at a lower translocation speed and with lower undulation frequency compared to two negative controls – the same animal under infrared light (black boxes) and the same strain not fed ATR (grey boxes); except for backward-crawling undulation frequency that was not statistically different under lime-colored and infrared light. Boxplots similar to Figure 3. ANOVA with Tukey pairwise comparison to the animal fed with ATR and under lime-colored light, n.s. p>0.05, ** p<0.1, **** p<0.0001. See Table 5-1 for details.

During free crawling or swimming, *C. elegans* exhibits a variety of behaviors from forward or backward almost-sinusoidal undulations, through omega and delta turns, direction changes, and pauses, to periods of quiescence (Stephens *et al*., 2008). When GABA transmission is impaired by knockout mutations or optogenetic inactivation, *C. elegans* becomes less coordinated, and their undulation frequencies and translocation speed decrease. In order to compare the efficiency of harnessing undulation frequency to produce translocation speed in wild-type and mutant animals, we isolated and analyzed the almost-sinusoidal undulations. For all strains, in both directions of crawling and swimming, there were positive linear correlations between undulation frequency and translocation speed. Wild type translocated faster with the same undulation frequency compared to any of the GABA transmission knockout strains (Fig. 6ab).

**Figure 6.**
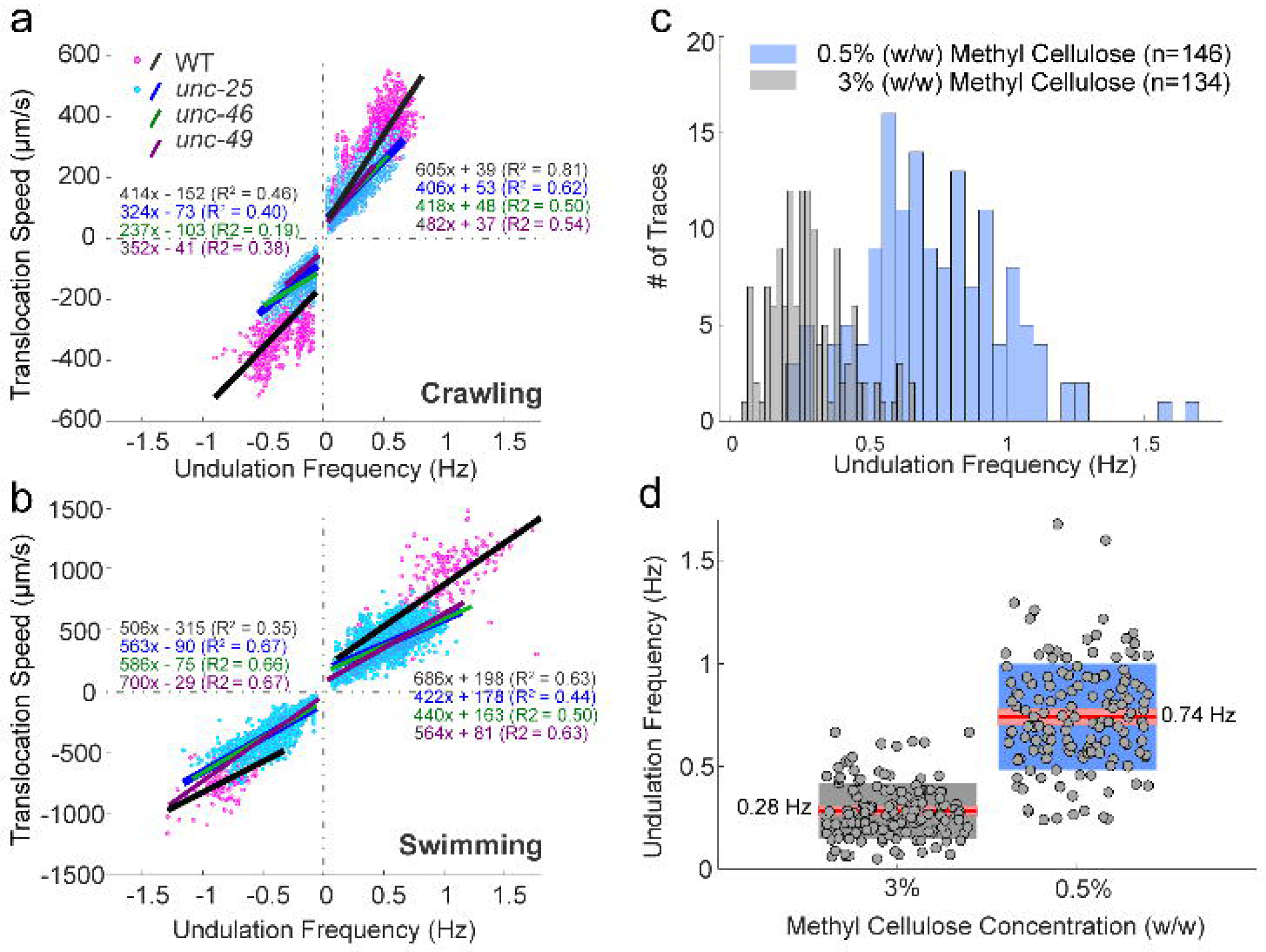
Undulation frequency correlated with translocation speed during free locomotion and undulations frequencies vary with the ambient viscosity in microfluidic waveform channels. (a, b) Undulation frequency was positively correlated with translocation speed during crawling and swimming. Wild type (magenta dots) and *unc-25* (cyan dots) data are during episodes in which animals performed sinusoidal locomotion. Fitted linear regression lines of wild type (black), *unc-25* (blue), *unc-46* (green) and *unc-49* (purple) strains; coefficients of determination (R^2^) and equations are labeled with the same colors. (c) In waveform microfluidic devices, the animals can undulate in predetermined channels designed to restrict their path with corresponding sequences of postures. When the ambient viscosity was high (3% w/w methyl cellulose solution) wild-type animals undulated with lower undulation frequencies than in less viscous environment, (0.5% w/w methyl cellulose solution). Frequency was measured by tracking the motion of neurons and muscle cells during calcium imaging. Sample size (n) is the number of cells analyzed. (d) Same data as c. The average undulation frequency in 3% methyl cellulose solution was 0.28 ± 0.14, and that in 0.5% solution was 0.74 ± 0.26 Hz. Red lines: mean values; pink boxes: 95% confidence intervals for the mean; gray and blue boxes: standard deviation; and gray dots: individual data points.

### GABAergic motoneurons showed different activation patterns during low and high undulation frequencies

In the absence of GABA transmission, animals did not move at rapid undulation frequency and fast translocation speed in either direction during free locomotion or in response to harsh touch stimulation to the head or tail. However, they were capable of slow dorsoventral undulation in both directions during free locomotion and after stimulation. Moreover, when only sinusoidal undulations are considered, undulation frequency is a predictor of translocation speed in wild-type and mutant animals, with wild type exhibiting higher efficiency. Together, these results suggest that GABAergic motoneurons’ contributions depend on undulation frequency and that they are necessary for high-frequency undulation.

To determine the contributions of GABAergic motoneurons during low and high undulation frequency, we recorded their activity with a genetically encoded calcium sensor. To keep track of the locomotive phase and undulation frequency we used a silicon microfluidic device with sinusoidal waveform channels designed to mimic the crawling path of wild-type animals (Fig. 7a; Lockery *et al*., 2008). Taking advantage of the predetermined shape of the channel and the published perimotor location of each motoneuron (Chen *et al*., 2006; Haspel and O’Donovan, 2011), we converted the soma position of GABAergic motoneurons during imaging to the locomotive phase of their neuromuscular junction to body-wall muscle. We assigned phases of the locomotion cycle to location in the channel according to the animal’s dorsoventral bending and direction of movement. We define the peak of dorsal bending as 90° and that of ventral bending as 270°, with other positions in the channel assigned accordingly. To manipulate the undulation frequency, we changed the viscosity of the fluid inside the channels with increasing concentration of methyl cellulose (in NGM buffer) up to 3% (w/w) methyl cellulose (higher concentrations prevented animals from entering the channels). When animals crawled in 3% (w/w) methyl cellulose, the mean undulation frequency was 0.28 ± 0.14 Hz, which is close to the undulation frequencies of the GABA transmission knockouts during forward crawling (0.32 ± 0.18 Hz, p = 0.89). Animals crawled faster in 0.5% (w/w) methyl cellulose, with a mean undulation frequency of 0.74 ± 0.26 Hz (Fig. 6cd). We imaged calcium level changes in the cytoplasm of body-wall muscle cells with GCaMP2 (TOL15) in 1.5% (w/w) methyl cellulose and in the cytoplasm of GABAergic motoneurons with GCaMP6 (TOL11) in 0.5% and 3% (w/w) methyl cellulose. The average undulation frequency in 1.5% methyl cellulose was 0.62 ± 0.20 Hz. Hence, regardless of methyl cellulose concentration, we used 0.6 Hz as the cut-off to categorize data into low and high undulation frequencies. Because the animals rarely moved backward in the channels, we only collected and analyzed calcium imaging data for smooth forward locomotion.

**Figure 7.**
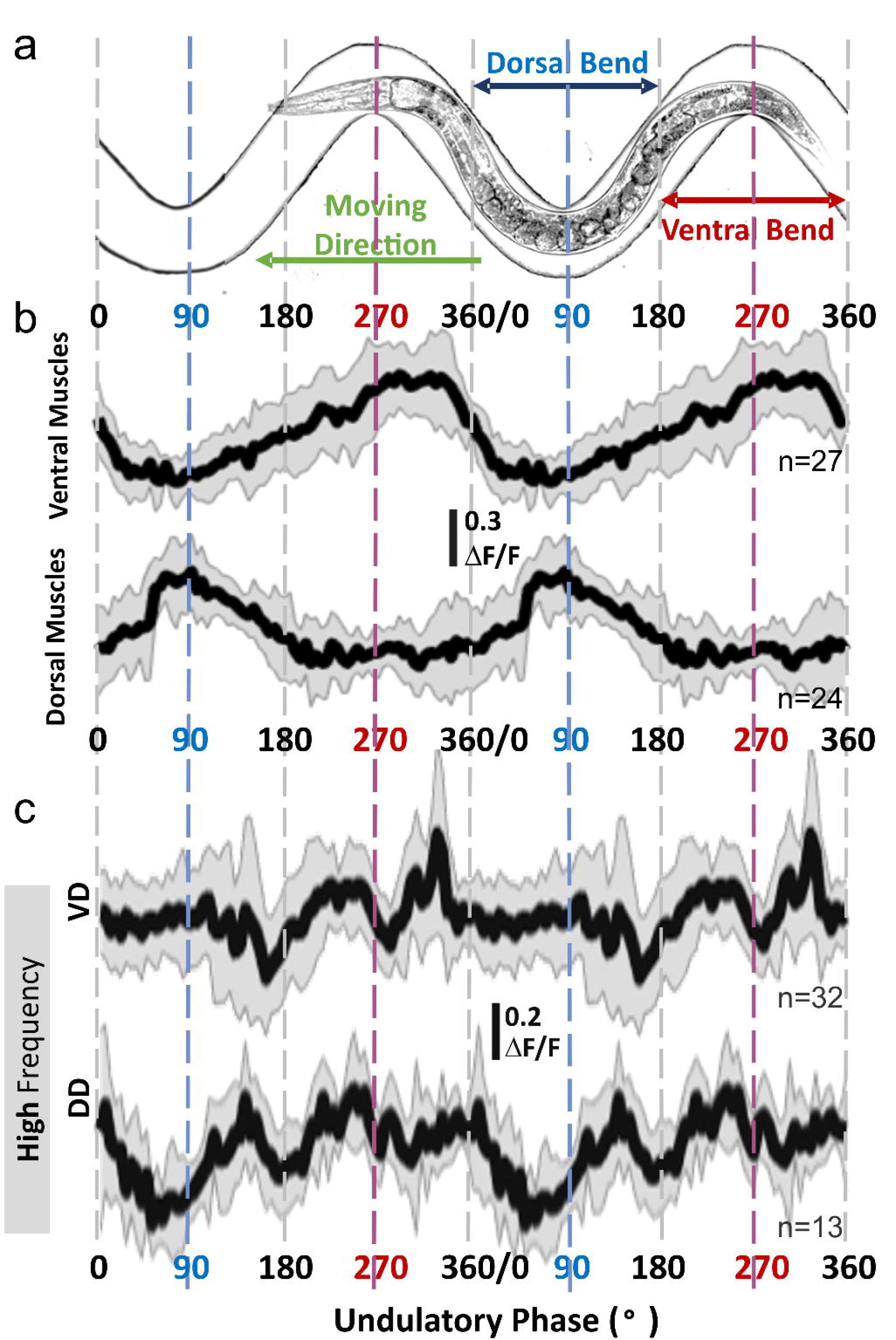
Calcium signal from GABAergic motoneurons relate to locomotive phase and muscle activity only during high undulation frequency. (a) Animals were free to move in sinusoidal microfluidic channels so that the phase of their undulatory cycle, including dorsal and ventral bends, is fixed and imposed by their spatial location. (b) Dorsal and ventral body-wall muscle cells were active during bending of the same side of the body (e.g., dorsal muscles were active during dorsal bend around 90°) regardless of undulation frequency. (c) During crawling at high undulation frequency, VD showed two troughs around 180° and 270°, and DD around 90° and 180°; only VD showed a sharp increase around 300°. The sample size (n) indicates the number of analyzed cells; solid lines and shaded areas in calcium traces are the mean and standard deviation. ΔF/F = (F_top50%_-F_BG_) / (∑(F_top50%_-F_BG_)/n). Changes in fluorescence levels and the perimotor-corrected locomotor phase were analyzed and calculated with a customized Matlab program (code 7-1).

The phase of the calcium imaging signal from body-wall muscle cells correlated to muscle contraction and bending. The signal increased in ventral muscle cells during ventral bending (around 270°, Fig. 7b) and in dorsal muscle cells during dorsal bending (around 90°), comparable to published calcium imaging signal during crawling on an agar surface (Butler *et al*., 2015). During high undulation frequency, GABAergic motoneurons exhibited a sustained level of calcium signal with distinct troughs of phasic lower levels (Fig. 7c). VD motoneurons exhibited phasic low signal around ventral bending (270°), while DD motoneurons exhibited a corresponding phase of low signal around dorsal bending (90°). Both VD and DD had a phase of low signal around 180°, and only VD had a rise in calcium signal level between 270° and 360°. In contrast, the calcium signal pattern in VD and DD during low undulation frequency were similar to each other and not different from a recording from calcium-insensitive green fluorescent protein (GFP) expressed in GABAergic motoneurons: a wide increase in signal from 180° to 360° that correlates with ventral bending. The increase in fluorescence intensity is likely due to convergence of fluorescence from neighboring motoneurons when the ventral nerve cord went through ventral bends.

Therefore, GABAergic motoneurons exhibited different patterns of activation during high and low undulation frequencies. In the high-frequency mode, VD and DD motoneurons have different inactive phases; in the low-frequency mode, VD and DD motoneurons showed similar wide inactive phases.

### Computational models tested three hypotheses for the role of inhibition in fast locomotion

Based on the experimental results, we suggest five hypotheses to explain the role of inhibition in sustaining high-frequency locomotion. First, it has long been suggested that cross-inhibition of the opposing body wall muscles increases the dorsoventral difference in muscle activation (McIntire 1993a, Bryden and Cohen 2008). Second, disinhibition of the innervated body wall muscles increases muscle activation, particularly during the rising phase. Third, inhibitory reset of VA and VB by the VD motoneurons allows higher locomotion frequency through phasic inhibition of ventral motoneurons (consistent with Boyle *et al*., 2012). Fourth, VD disinhibition of the VA and VB ventral cholinergic motoneurons amplifies, particularly during their rising phase. Fifth, reciprocal inhibition between VD and DD motoneurons stabilizes dorsoventral alternation.

From the experimental results, we found that *vab-7* mutants (in which DB neurons differentiate as VA) and *unc-4* mutants (in which DA and VA differentiate as DB and VB) were not different from the wild type in their shrinking response to harsh head and tail touch, respectively. This suggests that the mechanism for the shrinking response is not caused by coactivation of forward and backward premotor interneurons (AVA and AVB); instead, it appears localized within the motor circuit. Furthermore, shrinking appears qualitatively similar in the forward and backward motor circuits. We therefore focused our computational models on forward locomotion. Our model results can be reproduced for backward locomotion by changing the direction of the propagating wave under feedforward control or by changing the direction of the proprioceptive field under feedback control (Denham *et al*., 2018). Our model results are only applied within the scope of the function of inhibition, and we do not assume that forward and backward circuits are driven by the same mechanism nor do we model the transitions between forward and backward locomotion.

There are two prevalent hypotheses to explain the undulatory body bends in *C. elegans*: the proprioception (feedback-driven) hypothesis and the endogenous oscillator (feedforward-driven or central pattern generator) hypothesis. Proprioception transduces body curvature to adjacent body segments, so that rhythmic movement propagates along the body; endogenous oscillators (CPG circuits) generate rhythmic movement at the head or tail of the animal, or along the body (Gjorgjieva *et al*., 2014). Although proprioception and endogenous oscillators have been evidenced in *C. elegans*, their relative contributions during locomotion remain to be determined (Fouad *et al*., 2018; Gao *et al*., 2018; Xu *et al*., 2018). Previous models (Denham *et al*., 2018; Izquierdo and Beer, 2018) have investigated these regimes separately in order to parse their qualitative differences.

The models we used in this study are based on evidence for proprioceptive mechanisms (Wen *et al*., 2012) and for endogenous oscillators that could provide feedforward motor control (Gao *et al*., 2018). Using a proprioceptive model, we tested cross-inhibition, disinhibition, and inhibitory reset. Using a CPG-driven model, we tested muscle cross-inhibition and muscle disinhibition. The role of GABAergic motoneurons in rhythm generation in the *C. elegans* nerve cord remains uncharacterized and is not addressed in our models. In particular, more detailed neural models would be required to test VD disinhibition of VA and VB motoneurons and reciprocal inhibition between VD and DD motoneurons (Fig. 8a). In addition, a consequence of modeling purely feedforward, CPG-driven motor activity is the absence of a mechanism for the modulation of frequency, e.g., in response to changes in environmental viscosity. For both models, we simulated locomotion in two environments (agar-like and liquid-like) but did not directly manipulate undulation frequency within a given environment. We simulated forward locomotion in agar-like and liquid-like environments under proprioceptive and CPG control. Then, we compared simulations of wild-type animals with simulations that incorporate perturbations of the neuromuscular input due to the omission of cross-inhibition, disinhibition, or both (Fig. 8b).

**Figure 8.**
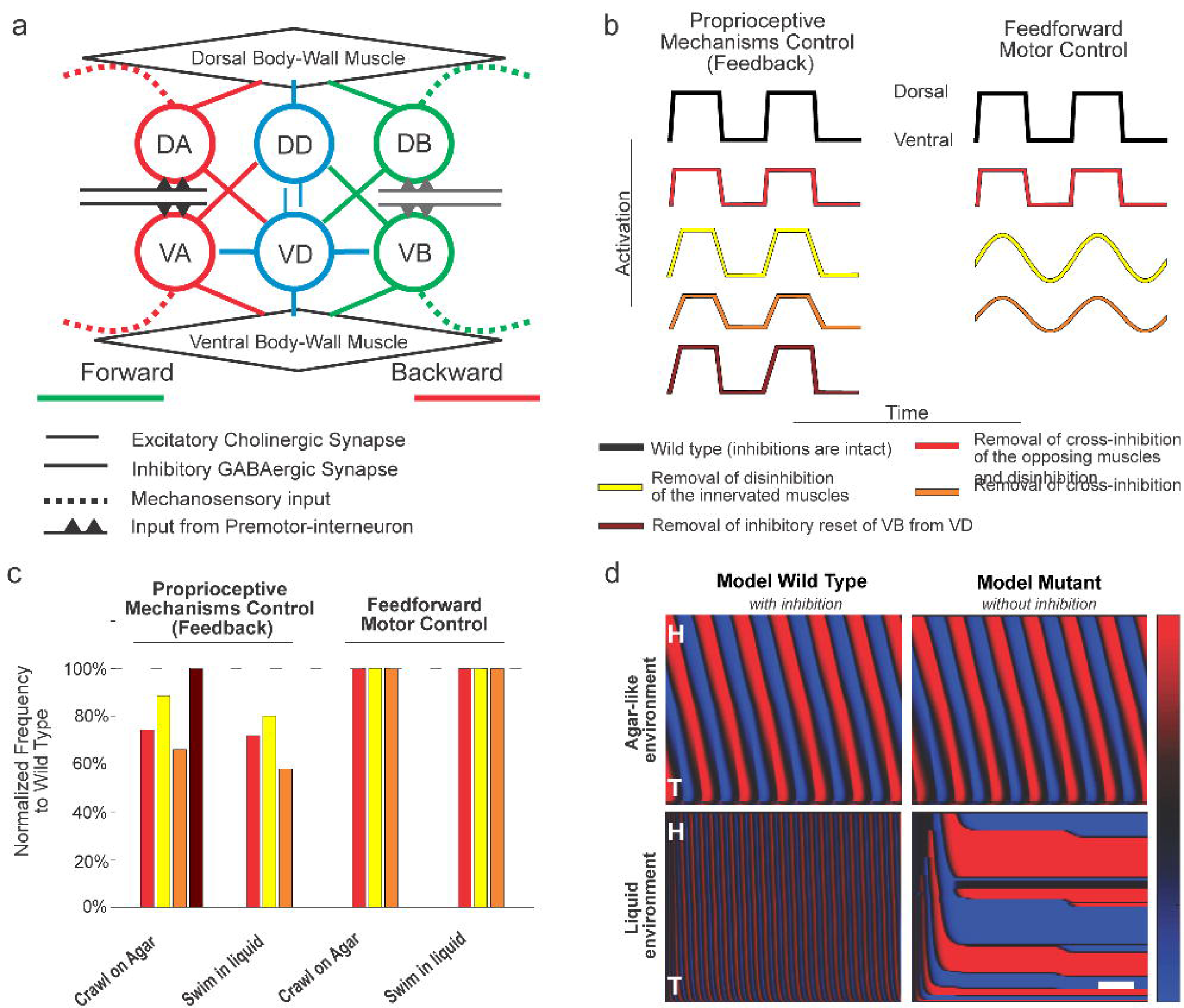
Integrated neuromehanical computational models test the role of inhibition in proprioceptive (feedback) driven pattern generation, as compared to CPG (purely feedforward) models. (a) Repeating units of the neuromuscular circuit were used to model *C. elegans* locomotion control. Single unit schematic motor circuit of the ventral nerve cord used in our proprioceptive model of locomotion. During forward locomotion, VB and VD (DB and DD) motoneurons innervate ventral (dorsal) muscles and VD motoneurons innervate VB motoneurons. During backward locomotion, A-type motoneurons take the place of B-type motoneurons. Postulated mechanosensation (dash lines) encode a proprioceptive signal by integrating the extent of bending over its receptive field. Suprathreshold proprioceptive currents trigger the activation of VB and DB motoneurons during forward locomotion (or VA and DA motoneurons during backward locomotion). The reciprocal synaptic connections between VD and DD motoneurons (blue thin lines) are not considered in the model. Adapted from Figure 1b of Cohen and Denham (2019). (b) Schematic traces of muscle activation in the proprioceptive control model (left) and feedforward control model (right), under different model perturbations. Positive activation denotes dorsal muscle activation. By construction, dorsal and ventral activation are in exact antiphase. Three hypotheses for the role of inhibition (cross-inhibition of the opposing muscles; disinhibition of the innervated muscles; and inhibitory reset of VB by VD) were tested in proprioceptive and feedforward motor control models. When muscle inhibition is disrupted, the amplitude or the shape of muscle input waveform is modified from the model wild type (black). Shown are schematics of perturbations corresponding to hypothesis 1 (reduced amplitude, red), hypothesis 2 (slower/smoother waveform, yellow), hypotheses 1 and 2 combined (orange), and hypothesis 3 (removal of neural inhibition, brown, proprioceptive model only). (c) Relative frequency changes due to model perturbations. Under models of proprioceptive control the undulation frequency is reduced by about 10% to 35% relative to the wild type under a single perturbation to muscle inhibition, or up to about 40% under the combined perturbation, suggesting that sufficient muscle activation amplitude and rapid activation onset are both required to sustain rapid undulations, during both crawling and swimming. Under proprioceptively driven locomotion, VB inhibition by VD serves to reset the neural rhythm during rapid undulations, but has negligible effect in slow crawling like motion. Bar plots with color scheme as in b depict the undulation frequency of model mutants normalized by the respective frequency of the model wild type. (d) Under a model of proprioceptive control, inhibitory reset is necessary for rapid swimming in a liquid environment (bottom), but not for slow crawling on agar-like environments (top). In an agar-like environment, the elimination of VD-to-VB inhibition (right) has no effect. In a water-like environment, elimination of this neural inhibition eliminates the rhythmic pattern altogether. Red and blue shaded areas represent dorsoventral curvature (vertical axis is the length of animal from the head, H, on the top to the tail, T; color bar is ventral (blue) to dorsal (red) curvature -10 to 10 mm^-1^), along time (horizontal axis, common scale bar is 1 s).

Under proprioceptive control, we find that all three perturbations induced a substantial decrease in undulation frequency (Fig. 8cd, Ext. Fig. 8-1). Simulated mutants lacking cross-inhibition of the opposing muscles showed the largest reduction (25% and 28% of simulated wild-type frequency in agar-like and liquid-like environments, respectively, in response to a 20% change in muscle input) followed by simulated mutants lacking disinhibitory effect on the innervated muscle (22% and 20% of wild-type frequency in agar-like and liquid-like environments, respectively, in response to a 20% change in muscle timescale). Applying both manipulations simultaneously decreased undulation frequency of the wild type even further (by 34% and 42% in agar-like and liquid-like environments, respectively). Intuitively, for proprioceptive control, continuous activation within the antagonistic muscle during bending causes stiffness of muscles, slowing down bending, and in turn delaying the state switch of the adjacent motoneurons (i.e., the propagation of the undulatory wave). The same simulations also showed a decrease in overall locomotion speed, which was proportional to the frequency decrease in liquid, but disproportionately small in agar. In contrast, in our model of CPG control, muscle inhibition had no effect on the undulation frequency (Fig. 8cd, Ext. Fig. 8-1). We also found that removing cross-inhibition of model muscles results in reduced speed, whereas removing muscle disinhibition resulted in increased locomotion speeds, indicating that the propagation of smooth sinusoidal, rather than square, muscle activation wave is mechanically more efficient (Lighthill, 1960). Together, these results suggest that the observed frequency– speed dependence is a signature of proprioceptively driven neural control. Our model further suggests that inhibition of GABAergic motoneurons influences undulation frequency primarily at the neuromuscular junctions. We note that additional frequency modulation mechanisms, which lie outside the scope of our model, cannot be ruled out under combined CPG and proprioceptive control.

We next tested the inhibitory resetting of VB by VD motoneurons. A neuromechanical model by Boyle *et al*., (2012) suggests that as a consequence of bistability of the A- and B-type motoneurons (Liu *et al*., 2014), ventral and dorsal motoneurons may simultaneously occupy the same state, effectively pausing locomotion in a manner that is reminiscent of the *shrinker* phenotype. To resume healthy locomotion, Boyle *et al*., (2012) proposed that inhibition from VD to VB motoneurons acts as a neural reset mechanism, switching VB off in order to facilitate a dorsal bend. Simulations of Boyle *et al*.’s model show that this neural reset has little effect on slow locomotion (in an agar-like environment) but is required for coordinated undulations during fast locomotion (in a liquid-like environment). This result inspired our third hypothesis and the reimplementation of this mechanism in the current model (see Methods). We simulated nematodes lacking neural inhibition from VD to VB motoneurons in our proprioceptive model. The simulated mutant did not show any significant change in frequency in an agar-like environment (Fig. 8cd). However, when simulated in a liquid-like environment, these simulated mutants became severely uncoordinated and failed to make progress, consistent with Boyle *et al*. (2012). This result indicates a requirement for a neural inhibitory reset mechanism to sustain coordinated fast undulations, at least in the absence of additional backup mechanisms (such as CPG control).

To summarize, our model results suggest that under proprioceptively driven locomotion, muscle cross-inhibition enhances locomotion frequency and speed. The associated frequency– speed dependence is consistent with our experimental observations. In this model, cross-inhibition and disinhibition of the body-wall muscles supports high-frequency undulatory locomotion, whereas the neural inhibition within the motor circuit can stabilize high-frequency undulations.

## Discussion

We investigated the role of inhibition in locomotion by analyzing the behavior of wild-type and mutant animals, recording neuronal and muscular activity in undulating intact animals, and by testing the role of inhibition under different control paradigms with computational models. Both wild type and GABA transmission mutants responded to harsh touch to the head or tail by shrinking, suggesting that dorsoventral coactivation is produced by the unimpaired nervous system. Impairment of GABA transmission genetically or optogenetically induced lower undulation frequency and translocation speed during crawling and swimming in both directions. GABAergic motoneurons’ activity patterns were different during undulation at low and high frequencies. During low undulation frequency, VD and DD showed a similar activity pattern, while during high undulation frequency, their activity alternated. We suggested five hypotheses for inhibitory mechanisms that support rapid undulation and tested three of them computationally. Our modeling results suggested that cross-inhibition and disinhibition of the body-wall muscles support rapid undulation. Finally, we suggest that the unimpaired locomotion circuit operates at two distinct modes to produce an undulatory motor program at low and high frequencies.

### Shrinking occurs in wild type and GABA transmission mutants

The shrinking response, presumably from coactivation of the dorsoventral muscle, has been described in GABA transmission mutants (McIntire *et al*., 1993a; McIntire *et al*., 1993b) and following laser ablation of GABAergic motoneurons (Yanik *et al*., 2004). Here, we show that shrinking commonly occurs in wild type in response to harsh touch stimuli to the head or tail. In particular, the tail stimuli induces a rapid forward movement with change in frequency and speed but not direction. Furthermore, the shrinking response does not require the simultaneous contribution of cholinergic motoneurons associated with the forward and backward directions. Neurodevelopmental mutant animals (namely, *vab-7* and *unc-4*) with impaired A- or B-type cholinergic motoneurons, respectively, show a shrinking response after the harsh touch stimulation that induces their normal undulatory locomotion (e.g., harsh head touch to *vab-7* induces wild-type-like sinusoidal backward undulation). These results suggest that shrinking is not associated with backward locomotion or with a change in direction, but rather with the switch from slow to fast locomotion and that shrinking is part of the motor program produced by the unimpaired nervous system. The switch in locomotion speed could be a specific case of a more general phenomenon in which simultaneous contraction of antagonistic muscle occurs during the initiation of rapid behavior.

The shrinking phenotype was described in mutant animals (McIntire *et al*., 1993a; McIntire *et al*., 1993b), but to date, has not been reported in wild-type animals, probably because the latter resolves the coactivation and swiftly moves away from noxious stimuli. Video microscopy and frame-by-frame inspection are therefore required to capture the brief body shrinkage of wild-type animals. Morphological measurement of the change in body length before and after harsh touch stimulation revealed that every animal, either wild type or the GABA transmission knockouts, shrank. It is the longer lasting shrinkage and lower translocation speed of GABA transmission mutants, which makes the observation of the shrinking phenotype easier with the naked eye.

### The distinct phenotype of the GABA transmission mutants is slow swimming

GABA transmission mutants, colloquially named ‘*shrinkers’*, such as *unc-46* and *unc-47* mutant strains defective in GABA vesicular transporter and *unc-49* mutant strain defective in anionic GABA_A_ receptors, have been described as defective in backward locomotion because they fail to produce wild-type-like swift reversal following a harsh touch on the head (McIntire *et al*., 1993a; McIntire *et al*., 1993b; Schuske *et al*., 2004). A similar phenotype was described in animals after laser ablation (Yanik *et al*., 2004). However, we found that the shrinking phenotype is not related to the target of noxious stimuli or the direction of locomotion, as GABA transmission mutants shrink in response to harsh touch to the tail as well as the head. Moreover, wild-type animals exhibit a shrinking phenotype, although smaller in magnitude and quicker to resolve. When suspended in liquid, GABA transmission mutants are uncoordinated and swim slowly with lower undulation frequency and translocation speed compared to those of the wild type. They exhibit a similar but less prominent phenotype when crawling on agar plates. Hence, slow and uncoordinated swimming, rather than shrinking, is a distinct phenotype of GABA transmission mutants.

### GABA transmission is necessary for fast dorsoventral alternation

During free crawling or swimming in either direction, an acute or chronic absence of GABA transmission leads to lower undulation frequency and translocation speed. The effect is more severe during swimming, which is typically performed at higher undulation frequency. Even following noxious touch stimuli that induce a rapid escape response in the wild type, GABA transmission mutants do not increase their undulation frequency and translocation speed and are slower than those of the wild type. These results deem GABA transmission necessary for fast locomotion. Similarly to our results, optogenetic activation of dopaminergic neurons induced a decrease in undulation frequency during crawling and swimming (Vidal-Gadea *et al*., 2011). That effect is, however, context-dependent. Hence, decrease in efficacy of GABA transmission could only partially explain the decrease in undulation frequency following increase in dopamine and their effect could be mediated by different mechanisms.

The undulation frequency dictates translocation speed during sinusoidal crawling and swimming behaviors in wild-type and mutant animals. Correlations between undulation frequency and translocation speed were always positive. The slope of the linear regression was always steeper for the wild type, implying a more efficient translation of undulation frequency to translocation speed. This suggests a subtle contribution of inhibition to the motor program of slow locomotion because the data for mutant animals is of slower translocation speed. Shape related parameters such as amplitude and wavelength are weakly correlated to instantaneous translocation speed, although they may contribute to fast locomotion.

### GABAergic motoneurons exhibit dorsoventral alternating inactive phases that match the activation of their postsynaptic body-wall muscles only during high-frequency undulation

Morphologically, GABAergic motoneurons synapse to the muscle arms of body-wall muscles (White *et al*., 1976, 1986), and their inhibitory effect on body-wall muscles was confirmed by laser ablation, muscle electrophysiology during bath application of GABA, and activation GABAergic motoneurons optogenetically (McIntire *et al*., 1993b; Bamber *et al*., 2005; Gao and Zhen, 2011; Inoue *et al*., 2015). The neuronal activity of GABAergic motoneurons has not been reported and was presumed to be similar to that of its antagonistic muscle, as both are activated by the same excitatory motoneurons. However, when we recorded the changes in calcium levels in GABAergic motoneurons during undulations, we found that they exhibited a baseline level of calcium signal interspaced by phases of lower signal and that the pattern of activity was frequency dependent. In high-frequency undulation, when calcium signal level in ventral body-wall muscle cells increased during contraction around 270° of the locomotion cycle, the calcium signal in VD motoneurons decreased; and when the calcium signal level in dorsal body-wall muscle cells increased during contraction around 90°, the calcium signal in DD motoneurons decreased (Fig. 7). These are possibly inactivity windows that allow postsynaptic muscles to escape from inhibition and be activated more sharply, or they could amplify muscle activity through disinhibition. In contrast, during low-frequency undulation, VD and DD exhibit similar patterns of calcium signal changes with a wide trough from 90° to 270°, which includes portions of ventral and dorsal bending regions. Hence, our calcium imaging of GABAergic motoneurons further supports their involvement in high-frequency undulations as their activity patterns are different for low and high undulation frequencies.

### Other possible inhibitory roles of GABAergic motoneurons in high-frequency undulation

GABAergic motoneurons might play several inhibitory roles during high-frequency undulation. We tested three hypotheses for the neuronal mechanisms that underlie these roles using proprioceptive mechanisms control and feedforward motor control models: 1) Cross-inhibition, in which GABAergic motoneurons inhibit the body wall muscle opposing an activated muscle; 2) disinhibition of the innervated body wall muscles, in which the release from inhibition enhances muscle activation; and 3) inhibitory reset of VA and VB, in which VD input terminates the locomotion cycle early and allows higher locomotion frequency (Boyle *et al*., 2012; Cohen and Sanders, 2014; Cohen and Denham, 2019). Our model results suggest that cross-inhibition and disinhibition of the body-wall muscles supports high-frequency undulatory locomotion, whereas the neural inhibition within the motor circuit stabilizes such high-frequency undulations. Besides the aforementioned roles of inhibition, there are other possible roles that we did not test because of experimental or computational model limitations. First, VD disinhibition of VA and VB, in which the activation of ventral cholinergic motoneurons is amplified, particularly during their rising phase. Second, reciprocal or nonreciprocal inhibition between DD and VD motoneurons, which might reinforce the antiphasic activity of GABAergic motoneurons during rapid locomotion. These inhibition or disinhibition are not exclusive in sustaining the fast dorsoventral body-wall muscle alternation. Cross-inhibition and disinhibition of muscles fulfill the activation–relaxation alternation of dorsoventral muscles, and neural inhibition helps to stabilize the motor circuit.

### Two modes of locomotion

We have demonstrated that the inhibitory motoneurons of the locomotion circuit are necessary only for rapid undulations and that their activity patterns are different for rapid and slow undulations. The frequency-dependent activity pattern of inhibitory motoneurons, the brief shrinking behavior when initiating an escape response, and the absence of rapid undulations when inhibition is inactive suggests a role for inhibition in the unimpaired nervous system. In addition, the distributions of frequency and translocation speed during crawling in both directions in the wild type are bimodal. Together, these results suggest that the locomotion circuit produces an undulatory motor program by two distinct modes of operation: low-frequency undulations that do not require inhibition and high-frequency undulations that require inhibition and in which GABAergic motoneurons exhibit an alternating activity pattern.

### Fundamental role for inhibition in rapid locomotion

Involvement of neuronal inhibition in the control of frequency and speed of behavior seems to be a fundamental mechanistic principle among locomotor circuits. In *Drosophila* larva, a set of inhibitory local interneurons, termed period-positive median segmental interneurons (PMSIs) control the frequency and speed of peristaltic locomotion by limiting the duration of motoneuronal bursting (Kohsaka *et al*., 2014). In adult *Drosophila*, similar yet unidentified inhibitory premotor input to leg motoneurons is associated with walking speed and their inactivation reveals an underlying slower walking speed (Gowda *et al*., 2018). In contrast, a lesion of the crossed glycinergic fibers or blocking glycinergic transmission in the lamprey spinal cord increase fictive burst frequency (Grillner, 2003). In the *Xenopus* tadpole phasic inhibition from the inhibitory interneuron pools aIN and cIN promotes faster swimming while weakening inhibition from dINs slows swimming down (Li and Moult, 2012). In zebrafish commissural local (CoLos) interneurons are recruited only during a rapid escape behavior (Liao and Fetcho, 2008; Satou *et al*., 2009). The glycinergic CoLos receive gap junctions from the Mauthner cell axon and inhibit the contralateral motoneurons as well as contralateral CiD and CoLo interneurons, thus inhibiting the driving force for contralateral muscle and disinhibiting the ipsilateral neuronal populations (Satou *et al*., 2009). In addition, ventrally located commissural bifurcating longitudinal (CoBLs) interneurons are only active at higher swimming frequencies (Liao and Fetcho, 2008). In the mammalian spinal cord (mostly studied in the mouse and cat), V1 neurons, a class of local circuit inhibitory interneurons are involved in the regulation of leg motoneuron bursting and step cycle duration to control the stepping frequency and speed of walking (Gosgnach *et al*., 2006). Genetic ablation or silencing of V1 neurons induces a marked decrease in locomotor frequency with no apparent effect of flexor-extensor coordination, while blocking synaptic output of both V1 and V2b neuronal populations, flexor–extensor pattern becomes synchronous (Zhang *et al*., 2014). In contrast, V0_D_ inhibitory commissural interneurons coordinate left-right alternation used in slow (walk) rather than fast (trot) gaits (Bellardita and Kiehn, 2015; Kiehn, 2016). Finally, a truncation mutation in the transcription factor gene dmrt3 gives Icelandic horses their typical tölting gait (characterized by the legs on the same body side moving forwards or backwards in synchrony), as well as pacing and ambling gaits in other breeds (Andersson *et al*., 2012). The protein DMRT3 is expressed in interneurons in the spinal cord, including inhibitory commissural neurons, suggesting that they are involved in gait and locomotion speed.

The frequency-supporting inhibitory inputs are mediated in insects and vertebrates by premotor interneurons that we suggest are analogous to motoneurons in the ventral nerve cord of *C. elegans* (Haspel *et al*., 2020). In this framework, the five pairs of nematode locomotion interneurons are analogous to descending interneurons (e.g., corticospinal or reticulospinal), whereas nematode motoneurons are analogous to spinal interneurons, integrating sensory and descending inputs, while generating and coordinating motor programs via their interconnectivity. It follows that nematode muscle arms are analogous to spinal motoneurons and each nematode muscle cell can be considered a motor unit. Accordingly, some *C. elegans* motoneurons are dedicated to a direction of locomotion (Haspel *et al*., 2010), while the muscle arms serve as the “final common path” (Sherrington, 1906). Hence, further study of the involvement of inhibition in rapid locomotion in *C. elegans*, where a small number of identifiable neurons can be recorded and controlled noninvasively with optical and genetic tools in freely moving animals, is likely to provide useful comparative insight for understanding the functional role, and possibly the evolutionary origin, of premotor inhibition in locomotory circuitry at large.

