## Supplementary figures and images for "Inhibition underlies fast undulatory locomotion in *C. elegans*"

### Extended Figure 8-1

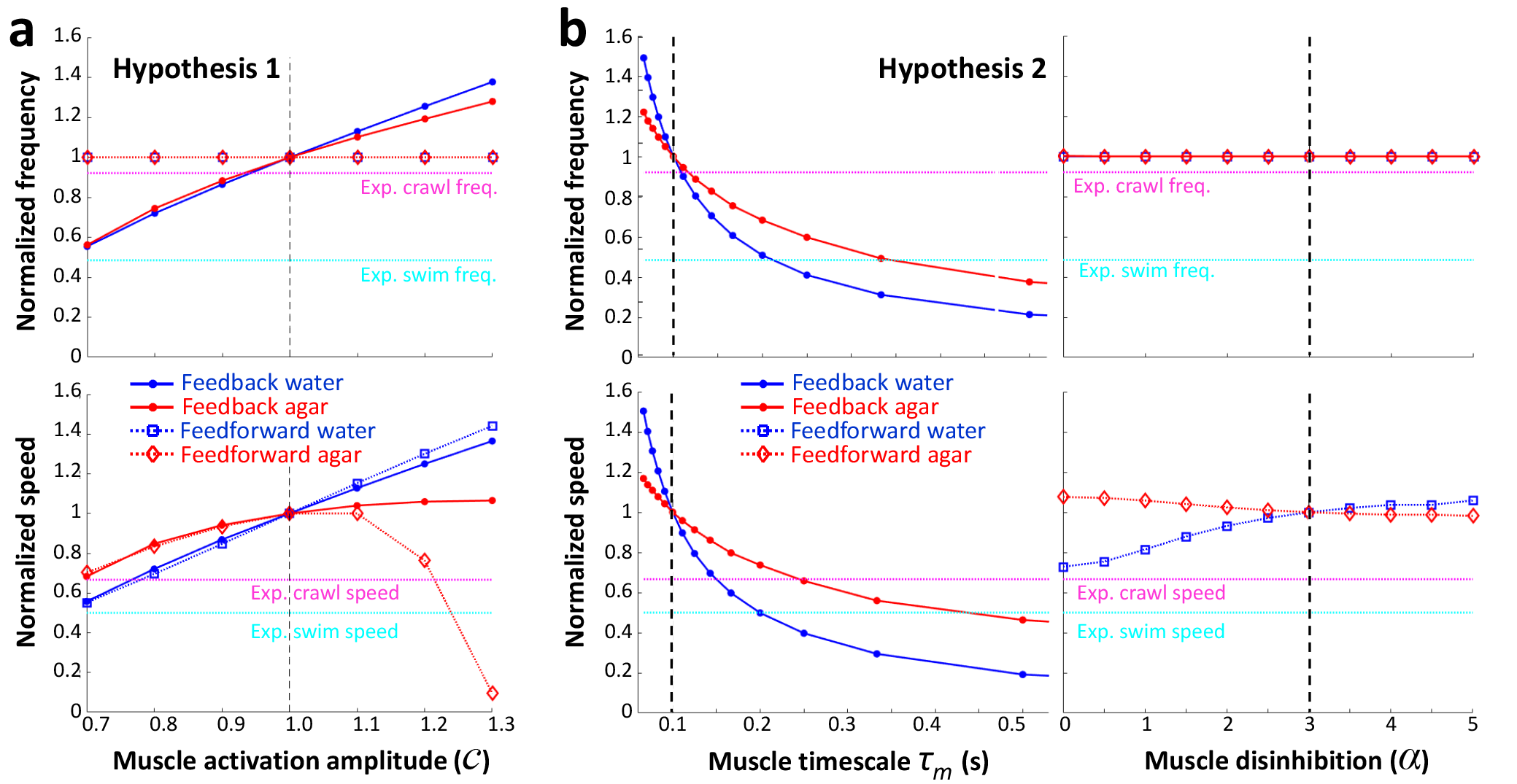
