## Supplemental table 2-2 for "Inhibition underlies fast undulatory locomotion in *C. elegans*"

**Table 2-2**. Wild-type animals move faster in response to head or tail touch while GABAergic mutant strains move slower or unchanged.

After either head or tail harsh touch, wild type animals (n=9) moved away at significantly higher translocation speed, undulation frequency and amplitude, while the mutant animals (n=9 for each strain) moved with unchanged or lower translocation speed and undulation frequency, compared to their locomotion before the stimulation. All the animals performed deeper bending with higher amplitude after stimulation while their wavelength did not change. Calculated probabilities for null hypotheses (p-value) below 0.05 were considered significant and are in red.

|  |  |  | **Head Stimulation** | | **Tail Stimulation** | |
| --- | --- | --- | --- | --- | --- | --- |
| **Wild Type** |  |  | Mean | p * | mean | p |
|  | Speed**  (µm/s) | Before | 199±51 | <0.0001 more | 244±76 | 0.0025 more |
|  |  | After | 345±51 |  | 366±62 |  |
|  | Frequency  (Hz) | Before | 0.39±0.07 | 0.0096 more | 0.44±0.12 | 0.0059 more |
|  |  | After | 0.45±0.06 |  | 0.58±0.08 |  |
|  | Amplitude  (µm) | Before | 183±30 | 0.0003 more | 184±35 | 0.0037 more |
|  |  | After | 323±41 |  | 260±60 |  |
|  | Wavelength  (µm) | Before | 662±109 | 0.1649 | 629±88 | 0.7068 |
|  |  | After | 630±50 |  | 615±46 |  |
| ***unc-25*** | Speed  (µm/s) | Before | 143±36 | 0.0170 less | 151±29 | 0.6912 |
|  |  | After | 76±48 |  | 131±101 |  |
|  | Frequency  (Hz) | Before | 0.33±0.07 | 0.0262 less | 0.36±0.07 | 0.1864 |
|  |  | After | 0.26±0.07 |  | 0.30±0.11 |  |
|  | Amplitude  (µm) | Before | 166±55 | 0.0462 more | 153±42 | 0.0008 more |
|  |  | After | 194±54 |  | 214±39 |  |
|  | Wavelength  (µm) | Before | 547±40 | 0.379 | 601±46 | 0.9813 |
|  |  | After | 605±202 |  | 601±54 |  |
| ***unc-46*** | Speed  (µm/s) | Before | 117±27 | 0.1578 | 123±28 | 0.0019 more |
|  |  | After | 54±96 |  | 171±44 |  |
|  | Frequency  (Hz) | Before | 0.32±0.13 | 0.0062 less | 0.38±0.13 | 0.0359 less |
|  |  | After | 0.20±0.09 |  | 0.30±0.09 |  |
|  | Amplitude  (µm) | Before | 174±46 | 0.0258 more | 167±33 | 0.0013 more |
|  |  | After | 224±53 |  | 229±39 |  |
|  | Wavelength  (µm) | Before | 668±80 | 0.0858 | 683±97 | 0.0107 less |
|  |  | After | 584±77 |  | 572±71 |  |
| ***unc-49*** | Speed  (µm/s) | Before | 140±28 | 0.0054 less | 139±15 | 0.8742 |
|  |  | After | 67±82 |  | 143±32 |  |
|  | Frequency  (Hz) | Before | 0.29±0.06 | 0.192 | 0.25±0.04 | 0.2661 |
|  |  | After | 0.25±0.13 |  | 0.33±0.06 |  |
|  | Amplitude  (µm) | Before | 176±41 | 0.1673 | 181±50 | 0.062 |
|  |  | After | 217±30 |  | 220±48 |  |
|  | Wavelength  (µm) | Before | 585±88 | 0.5487 | 562±47 | 0.9082 |
|  |  | After | 578±122 |  | 600±114 |  |
| * p value, paired two-tail T-Test | | | |  |  |  |
| **Speeds are the absolute values regardless of moving directions. | | | | | | |
