## Supplemental table 3-1 for "Inhibition underlies fast undulatory locomotion in *C. elegans*"

**Table 3-1**. During free crawling GABAergic mutants move slower than wild type and with smaller amplitude and wavelength.

Animals of three GABA transmission knockout strains crawled on agar surface with significantly lower translocation speed, undulation frequency, maximal amplitude, and primary wavelength. The only exception is that unc-25/GAD knockout animals move at the same mean frequency as wild type when moving forward. Calculated probabilities for null hypotheses (p-value) below 0.05 were considered significant and are in red.

|  |  | **Translocation Speed (µm/s)** | | | **Undulation Frequency (Hz)** | | | **Maximal Amplitude (µm)** | | | **Primary Wavelength (µm)** | | |
| --- | --- | --- | --- | --- | --- | --- | --- | --- | --- | --- | --- | --- | --- |
|  |  | **Mean±SD** | **One-way ANOVA** | **p value (Tukey test)** | **Mean±SD** | **One-way ANOVA** | **p value (Tukey test)** | **Mean±SD** | **One-way ANOVA** | **p value (Tukey test)** | **Mean±SD** | **One-way ANOVA** | **p value (Tukey test)** |
| **Backward Crawling** | **Wild Type** | 108±47 | F(3,878) = 64.73,  p < 0.0001 | Comparison Reference | 0.40±0.17 | F(3,431) = 8.65,  p < 0.0001 | Comparison Reference | 227±66 | F(3,878) = 37.22,  p < 0.0001 | Comparison Reference | 730±260 | F(3,843) = 16.42,  p < 0.0001 | Comparison Reference |
|  | ***unc-25*** | 70±28 |  | <0.0001 | 0.32±0.38 |  | 0.0295 | 168±57 |  | < 0.0001 | 644±113 |  | < 0.0001 |
|  | ***unc-46*** | 63±22 |  | <0.0001 | 0.32±0.25 |  | 0.0307 | 177±62 |  | < 0.0001 | 655±114 |  | < 0.0001 |
|  | ***unc-49*** | 34±22 |  | <0.0001 | 0.25±0.36 |  | <0.0001 | 208±64 |  | 0.0069 | 646±156 |  | < 0.0001 |
| **Forward Crawling** | **Wild Type** | 148±114 | F(3,987) = 136.49,  p < 0.0001 | Comparison Reference | 0.37±0.2 | F(3,682) = 14.10,  p < 0.0001 | Comparison Reference | 225±55 | F(3,987) = 44.01,  p < 0.0001 | Comparison Reference | 716±125 | F(3,955) = 10.98,  p < 0.0001 | Comparison Reference |
|  | ***unc-25*** | 90 ±69 |  | <0.0001 | 0.37±0.17 |  | 0.9997 | 176±50 |  | < 0.0001 | 662±90 |  | < 0.0001 |
|  | ***unc-46*** | 73±69 |  | <0.0001 | 0.31±0.19 |  | 0.0314 | 181±55 |  | < 0.0001 | 668±99 |  | < 0.0001 |
|  | ***unc-49*** | 47±67 |  | <0.0001 | 0.25±0.16 |  | <0.0001 | 212±60 |  | 0.0461 | 665±143 |  | < 0.0001 |
