## Supplemental table 4-1 for "Inhibition underlies fast undulatory locomotion in *C. elegans*"

**Table 4-1**. During free swimming GABAergic mutants move slower than wild type and with smaller amplitude and wavelength.

Animals of three GABA transmission knockout strains swam in saline with significantly lower translocation speed, undulation frequency, maximal amplitude, and primary wavelength than wild type. Calculated probabilities for null hypotheses (p-value) below 0.05 were considered significant and are in red.

|  |  | **Translocation Speed (µm/s)** | | | **Undulation Frequency (Hz)** | | | **Maximal Amplitude (µm)** | | | **Primary Wavelength (µm)** | | |
| --- | --- | --- | --- | --- | --- | --- | --- | --- | --- | --- | --- | --- | --- |
|  |  | **Mean±SD** | **One-way ANOVA** | **p value (Tukey test)** | **Mean±SD** | **One-way ANOVA** | **p value (Tukey test)** | **Mean±SD** | **One-way ANOVA** | **p value (Tukey test)** | **Mean±SD** | **One-way ANOVA** | **p value (Tukey test)** |
| **Backward Swimming** | **Wild Type** | -367±145 | F(3,2870) = 802.42  p < 0.0001 | Comparison Reference | 1.26±0.49 | F(3,2256) = 373.06  p < 0.0001 | Comparison Reference | 294±70 | F(3,2863) = 430.86  p < 0.0001 | Comparison Reference | 866±91 | F(3,2715) = 316.55  p < 0.0001 | Comparison Reference |
|  | ***unc-25*** | -165±42 |  | <0.0001 | 0.75±0.49 |  | <0.0001 | 209±52 |  | < 0.0001 | 691±84 |  | < 0.0001 |
|  | ***unc-46*** | -150±30 |  | <0.0001 | 0.92±0.31 |  | <0.0001 | 186±49 |  | < 0.0001 | 746±59 |  | < 0.0001 |
|  | ***unc-49*** | -84±18 |  | <0.0001 | 0.43±0.27 |  | <0.0001 | 230±47 |  | < 0.0001 | 729±125 |  | < 0.0001 |
| **Forward Swimming** | **Wild Type** | 383±132 | F(3,3163) = 1391.20  p < 0.0001 | Comparison Reference | 1.65±0.26 | F(3,2788) = 1684.89  p < 0.0001 | Comparison Reference | 273±38 | F(3,3163) =927.68  p < 0.0001 | Comparison Reference | 885±59 | F(3,3082) = 830.09  p < 0.0001 | Comparison Reference |
|  | ***unc-25*** | 188 ±77 |  | <0.0001 | 0.81±0.27 |  | <0.0001 | 194±39 |  | < 0.0001 | 694±73 |  | < 0.0001 |
|  | ***unc-46*** | 168±53 |  | <0.0001 | 0.95±0.27 |  | <0.0001 | 178±34 |  | < 0.0001 | 743±50 |  | < 0.0001 |
|  | ***unc-49*** | 94±51 |  | <0.0001 | 0.45±0.33 |  | <0.0001 | 225±44 |  | < 0.0001 | 738±123 |  | < 0.0001 |
