## Supplemental table 5-1 for "Inhibition underlies fast undulatory locomotion in *C. elegans*"

**Table 5-1.** Optogenetic inactivation of all GABAergic neurons during free crawling and swimming reduces locomotion speed.

When GABAergic motoneurons were acutely inactivated (animals fed with all trans retinal (ATR) and illuminated with lime-colored light), they crawled on agar surface or swam in saline with significantly lower translocation speed and undulation frequency. They were compared to two negative controls -- the same animal under infrared light (ATR / Infrared Light) and the same strain not fed ATR (No ATR / Infrared Light). The only exception is that backward-crawling undulation frequency (but not translocation speed) of animals that were fed ATR were not statistically different under lime-colored or infrared light. Calculated probabilities for null hypotheses (p-value) below 0.05 were considered significant and are in red.

|  |  | | **Translocation Speed (µm/s)** | | | **Undulation Frequency (Hz)** | | |
| --- | --- | --- | --- | --- | --- | --- | --- | --- |
| **Locomotion** | **Conditions** | | Mean±SD | One-way ANOVA | p value (Tukey test) | Mean±SD | One-way ANOVA | p value (Tukey test) |
| **Backward Crawling** | **Control** | **No ATR /**  **Infrared Light** | -123±106 | F(2,397) = 16.48,  p < 0.0001 | <0.0001 | 0.42±0.18 | F(2,195) = 5.03,  p = 0.0074 | 0.0074 |
|  |  | **ATR /**  **Infrared Light** | -101±81 |  | 0.0017 | 0.34±0.18 |  | 0.7423 |
|  | **Experimental** | **ATR /**  **Lime Light** | -67±52 |  | Comparison Reference | 0.32±0.18 |  | Comparison Reference |
| **Forward Crawling** | **Control** | **No ATR /**  **Infrared Light** | 201±72 | F(2,918) = 83.66,  p < 0.0001 | <0.0001 | 0.40±0.09 | F(2,845) =20.16,  p < 0.0001 | <0.0001 |
|  |  | **ATR /**  **Infrared Light** | 171±66 |  | <0.0001 | 0.37±0.09 |  | 0.0079 |
|  | **Experimental** | **ATR /**  **Lime Light** | 133±68 |  | Comparison Reference | 0.34±0.10 |  | Comparison Reference |
| **Backward Swimming** | **Control** | **No ATR /**  **Infrared Light** | -312±103 | F(2,783) = 111.59,  p < 0.0001 | <0.0001 | 1.11±0.20 | F(2,617) = 47.63,  p < 0.0001 | <0.0001 |
|  |  | **ATR /**  **Infrared Light** | -248±53 |  | <0.0001 | 1.01±0.39 |  | <0.0001 |
|  | **Experimental** | **ATR /**  **Lime Light** | -140±26 |  | Comparison Reference | 0.73±0.43 |  | Comparison Reference |
| **Forward Swimming** | **Control** | **No ATR /**  **Infrared Light** | 315±102 | F(2,899) = 192.43,  p < 0.0001 | <0.0001 | 1.36±0.30 | F(2,840) = 181.45,  p < 0.0001 | <0.0001 |
|  |  | **ATR /**  **Infrared Light** | 247±88 |  | <0.0001 | 1.07±0.28 |  | <0.0001 |
|  | **Experimental** | **ATR /**  **Lime Light** | 162±84 |  | Comparison Reference | 0.76±0.46 |  | Comparison Reference |
