## Supplementary material for "Inhibition underlies fast undulatory locomotion in *C. elegans*": Caption Extended Figure 8-1

**Extended Figure 8.** Suppression of muscle cross-inhibition in computational models revealed differences between putative roles of inhibition in CPG and proprioceptive driven control of locomotion. Normalized undulation frequency (top) and locomotion speed (bottom) plotted as a function of muscle parameters representing (a) muscle activation (testing Hypothesis 1) and (b) muscle waveform (testing Hypothesis 2). See Figure 8 and text for details regarding hypotheses. Undulation frequency and locomotion speed are normalized by the model output at wild-type parameter values (black vertical dashed lines) in agar-like (red) and water-like (blue) environments. Horizontal dashed lines show frequencies and locomotion speeds in GABA defective nematodes (*unc-25* experimental results) in agar (magenta) and water (cyan), normalized by the respective wild type values. Experimentally, both frequency and speed are reduced in GABA-defective animals as compared to the wild type with a relatively larger reduction in water than on agar. (a) Hypothesis 1: Muscle cross-inhibition enhances undulation frequency and locomotion speed. In the proprioceptive model, suppression of muscle cross-inhibition by approximately 8% and 30% quantitatively captures experimental drop in undulation frequency and speed on agar. Suppression of muscle cross-inhibition by the same amount in water results in a similar modulation of frequency. About 33% suppression of inhibition would be required to quantitatively account for the change in locomotion frequency in water, suggesting an additional roles of inhibition. Model speed changes with reduced muscle cross-inhibition ($C<1$) are similar in CPG and proprioceptively controlled models. In the CPG controlled model, the reduction in muscle amplitude leads to reduced locomotion speed, even at a fixed frequency, with a stronger speed reduction in water than in agar. (b) Hypothesis 2: Muscle disinhibition increases the muscle response rate. In a proprioceptive control model (left), removal of disinhibition is modeled by an increased muscle time scale. Muscle time scales ($\tau$) of 0.12 s and 0.25 s would account for the observed drop in frequency in water and agar, respectively; muscle time scales of 0.2 s would account for relative changes in model speeds. Under models of fixed-frequency CPG control (right), removal of disinhibition is modeled by a change of the waveform of muscle activation (from a near-square wave of the model wild type at $\alpha=3$ to a near sinusoidal wave for $\alpha\ll1$, see methods and Fig. 8b). This change in muscle activation waveform results in opposite relative changes in speed on agar and in water: speed on agar changes only modestly and is higher for smoother waveforms; speed in water is up to about 20% faster for squarer waveforms.
