## Supplemental table 8-1 for "Inhibition underlies fast undulatory locomotion in *C. elegans*"

Table 8-1: Wild type model parameters.

|  | **Description** | **Parameter** | **Value** |
| --- | --- | --- | --- |
| **Body geometry** | Body length  Cuticle thickness  Maximum body radius | $L$  $r_{\mathrm{cuticle}}$  $R$ | 1 mm  0.5 μm  40 μm |
| **Muscles** | Muscle timescale (feedback model)  Muscle disinhibition (CPG model)  Curvature amplitude | $\tau_{m}$  $\alpha$  $\beta_{0}$ | 0.1 s  $3.0$  $10$ mm^-1^ |
| **Mechanics** | Young’s Modulus  Tangential drag coefficient  Normal drag coefficient | $E$  $K_{\tau,\mathrm{water}}$  $K_{\tau,\mathrm{agar}}$  $K_{v,\mathrm{water}}$  $K_{v,\mathrm{agar}}$ | $10^{5}$ Pa  $3.3$x$10^{-3}$kgm^-1^s^-1^ $3.2$ kgm^-1^s^-1^  $5.2$x$10^{-3}$kgm^-1^s^-1^  $128$ kgm^-1^s^-1^ |
| **CPG  (feedforward) control** | Undulation wavelength (water)  Undulation wavelength (agar)  Undulation period (water)  Undulation period (agar) | $\lambda_{f}$  $\lambda_{f}$  $T_{f}$  $T_{f}$ | $0.6$ mm  $1.6$mm  $0.6$ s  $2.0$ s |
| **Proprioceptive control** | Ventral ON/dorsal OFF thresholds  Ventral OFF/dorsal ON threshold  Proprioceptive range  VB, DB NMJ weight  VD, DD NMJ weight  VD → VB inhibitory weight | $\theta_{V, ON,},\theta_{D,OFF}$  $\theta_{V, OFF}\theta_{D,ON}$  $\delta$  ${w^{\mathrm{exc}}}_{\mathrm{NMJ}}$  ${w^{\mathrm{inh}}}_{\mathrm{NMJ}}$  $w$ | 3.0  $-$3.0  0.5  0.8  $0.2$  $0.2$ |
