## Supplemental table 2-1 for "Inhibition underlies fast undulatory locomotion in *C. elegans*"

**Table 2-1.** Harsh touch stimulation to the head or tail induces shortening of body length in wild-type and a larger shortening in GABAergic mutants.

Body length of wild type and GABA transmission knockout animals (*unc-25*, *unc-46*, *unc-47* and *unc-49*) reduced after harsh stimuli to the head or the tail, but length reduced less and recovered sooner in wild type animals. Percentage of the body length at four time points: right away (0 s), 0.5 s, 1 s, and 2 s , was compared to the pre-stimulus value. Statistical tests compared mutants to wild type (ANOVA and Tukey) as well as post to prestimulus values (paired T test). Calculated probabilities for null hypotheses (p-value) below 0.05 were considered significant and are in red.

|  |  | **0 s** | | | | **0.5 s** | | | | **1 s** | | | | **2 s** | | | |
| --- | --- | --- | --- | --- | --- | --- | --- | --- | --- | --- | --- | --- | --- | --- | --- | --- | --- |
|  |  | Mean ± SD | p (Paired T-test) | One-way ANOVA | p (Tukey test) | Mean ± SD | p (Paired T-test) | One-way ANOVA | p value (Tukey test) | Mean ± SD | p (Paired T-test) | One-way ANOVA | p (Tukey test) | Mean ± SD | p (Paired T-test) | One-way ANOVA | p (Tukey test) |
| **Head touch stimulation** | **Wild Type** | 94±2% | 0.0001 | F(4,40) = 11.2 p < 0.0001 | Comparison Reference | 96±2% | 0.002 | F(4,40) = 3.91 p =0.009 | Comparison Reference | 97±3% | 0.0208 | F(4,40) = 3.68 p =0.0121 | Comparison Reference | 98±4% | 0.141 | F(4,40) = 3.91 p =0.0091 | Comparison Reference |
|  | ***unc-25*** | 90±2% | < 0.0001 |  | 0.043 | 92±4% | 0.0003 |  | 0.421 | 94±4% | 0.002 |  | 0.733 | 94±5% | 0.004 |  | 0.417 |
|  | ***unc-46*** | 91±4% | < 0.0001 |  | 0.285 | 93±5% | 0.005 |  | 0.645 | 94±5% | 0.012 |  | 0.770 | 95±4% | 0.010 |  | 0.543 |
|  | ***unc-49*** | 90±2% | < 0.0001 |  | 0.020 | 90±5% | 0.0003 |  | 0.045 | 92±5% | 0.001 |  | 0.085 | 92±6% | 0.004 |  | 0.067 |
|  | ***vab-7*** | 97±3% | 0.209 |  | 0.124 | 97±4%, | 0.039 |  | 0.975 | 99±2%, | 0.083 |  | 0.874 | 99±2%, | 0.322 |  | 0.950 |
| **Tail touch stimulation** | **Wild Type** | 95±3% | 0.003 | F(4,40) = 3.73 p =0.0114 | Comparison Reference | 98±3% | 0.1107 | F(4,40) = 9.54 p<0.0001 | Comparison Reference | 99±3% | 0.214 | F(4,40) = 6.89  p =0.0003 | Comparison Reference | 99±2% | 0.263 | F(4,40) = 3.27 p =0.0207 | Comparison Reference |
|  | ***unc-25*** | 91±3% | < 0.0001 |  | 0.040 | 92±5% | 0.0008 |  | 0.030 | 95±5% | 0.011 |  | 0.324 | 97±4% | 0.066 |  | 0.801 |
|  | ***unc-46*** | 92±3% | < 0.0001 |  | 0.161 | 90±4% | 0.0001 |  | 0.005 | 92±4% | 0.0004 |  | 0.005 | 93±4% | 0.003 |  | 0.072 |
|  | ***unc-49*** | 91±4% | 0.0001 |  | 0.039 | 88±6% | 0.0003 |  | 0.0001 | 91±4% | 0.0005 |  | 0.003 | 94±5% | 0.005 |  | 0.032 |
|  | ***unc-4*** | 94±2% | 0.0002 |  | 0.963 | 97±4% | 0.060 |  | 0.999 | 98±3% | 0.076 |  | 0.998 | 97±3% | 0.018 |  | 0.892 |
| **50 mM CuSO_4_** | **Wild Type** | 97±1% | 0.0001 |  |  |  |  |  |  |  |  |  |  |  |  |  |  |
| **0.1% SDS** | **Wild Type** | 99±1% | 0.205 |  |  |  |  |  |  |  |  |  |  |  |  |  |  |
| **Head touch stimulation** | **Paralyzed Wild Type** | 100±1% | 0.106 |  |  |  |  |  |  |  |  |  |  |  |  |  |  |
