## Supplemental table 2-3 for "Inhibition underlies fast undulatory locomotion in *C. elegans*"

**Table 2-3.** GABAergic mutant strains move at lower speed and frequency than wild-type 5 seconds post-stimulus.

Five seconds after a harsh touch to the head or tail all GABAergic knockout animals moved with lower translocation speed and undulation frequency than wild type. Calculated probabilities for null hypotheses (p-value) below 0.05 were considered significant and are in red.

|  |  | **Translocation Speed (µm/s)*** | | | **Undulation Frequency (Hz)** | | |
| --- | --- | --- | --- | --- | --- | --- | --- |
|  |  | **Mean±SD** | **One-way ANOVA** | **p value (Tukey test)** | **Mean±SD** | **One-way ANOVA** | **p value (Tukey test)** |
| **Head Stimulation** | **Wild Type** | 345±51 | F(3,35)=67.78, p< 0.0001 | Comparison Reference | 0.45±0.06 | F(3,35)=13.70, p< 0.0001 | Comparison Reference |
|  | ***unc-25*** | 76±48 |  | < 0.0001 | 0.26±0.07 |  | 0.0004 |
|  | ***unc-46*** | 54±95 |  | < 0.0001 | 0.20±0.09 |  | <0.0001 |
|  | ***unc-47*** | 67±82 |  | < 0.0001 | 0.25±0.13 |  | 0.0002 |
| **Tail Stimulation** | **Wild Type** | 366±62 | F(3,35)=49.89, p< 0.0001 | Comparison Reference | 0.58±0.08 | F(3,35)=20.11, p< 0.0001 | Comparison Reference |
|  | ***unc-25*** | 131±101 |  | < 0.0001 | 0.30±0.11 |  | < 0.0001 |
|  | ***unc-46*** | 171±44 |  | < 0.0001 | 0.30±0.09 |  | < 0.0001 |
|  | ***unc-47*** | 143±32 |  | < 0.0001 | 0.33±0.06 |  | < 0.0001 |
| *Translocation speeds are the absolute values regardless of moving directions. | | | | | | | |
